# Movie-trained transformer reveals novel response properties to dynamic stimuli in mouse visual cortex

**DOI:** 10.1101/2025.09.16.676524

**Authors:** Bryan M. Li, Wolf De Wulf, Danai Katsanevaki, Arno Onken, Nathalie L. Rochefort

## Abstract

Understanding how the brain encodes complex, dynamic visual stimuli remains a fundamental challenge in neuroscience. Here, we introduce ViV1T, a transformer-based model trained on natural movies to predict neuronal responses in mouse primary visual cortex (V1). ViV1T outperformed state-of-the-art models in predicting responses to both natural and artificial dynamic stimuli, while requiring fewer parameters and reducing runtime. Despite being trained exclusively on natural movies, ViV1T accurately captured core V1 properties, including orientation and direction selectivity as well as contextual modulation, despite lacking explicit feedback mechanisms. ViV1T also revealed novel functional features. The model predicted a wider range of contextual responses when using natural and model-generated surround stimuli compared to traditional gratings, with novel model-generated dynamic stimuli eliciting maximal V1 responses. ViV1T also predicted that dynamic surrounds elicited stronger contextual modulation than static surrounds. Finally, the model identified a subpopulation of neurons that exhibit contrast-dependent surround modulation, switching their response to surround stimuli from inhibition to excitation when contrast decreases. These predictions were validated through semi-closed-loop *in vivo* recordings. Overall, ViV1T establishes a powerful, data-driven framework for understanding how brain sensory areas process dynamic visual information across space and time.

Code available at github.com/bryanlimy/ViV1T-closed-loop.

## 1 Introduction

Understanding how the visual system processes high-dimensional, natural visual stimuli remains a major challenge in neuroscience. Until recently, research in this field has been constrained by the limited computational tools to interpret neuronal responses to complex stimuli. The emergence of advanced machine learning techniques, along with large-scale *in vivo* neuronal recording technologies [76, 77], has opened new opportunities for analysing extensive datasets of natural stimuli and predicting neuronal responses to diverse combinations of these stimuli [11, 51, 64, 85]. Predictive models of neuronal responses to natural stimuli enable large-scale, unbiased exploration of visual stimulus spaces *in silico*, which would not be possible in the limited time of *in vivo* recordings. Model predictions can then be validated using *in vivo* neuronal recordings with a selective set of relevant visual stimuli.

As deep neural networks (DNNs) have gained popularity in computational neuroscience [9, 43, 65], they have established new benchmarks in prediction accuracy and proven effective for predicting responses in mouse visual cortex [4, 27, 29, 30, 44, 48, 91]. DNN-based data-driven models share a common representation by being trained end-to-end directly on data from thousands of neurons, without any assumption on their connectivity. These models typically use a core-readout architecture, where a shared core captures a latent representation of visual stimuli across animals, and animal-specific linear readouts map these features to specific neurons [16, 18, 30, 44, 45, 48]. While convolutional neural networks (CNNs) are commonly used as cores, transformer models have recently gained traction [8, 45]. A key advantage of transformer-based models lies in their high performance in capturing dynamic processes [88]: their self-attention mechanisms can capture longrange dependencies, which may better reflect temporal responses of neurons to dynamic visual stimuli.

In natural environments, animals must rapidly interpret dynamic visual scenes to detect motion, avoid predators, or pursue prey. Despite the key role of visual cortex in motion perception [49], previous work has focused on static predictive models of mouse visual cortex (*i.e*. responses to static images) [31, 45, 48, 85]. These predictions included the generation of most-exciting images (MEIs) that were optimised to elicit maximal responses of single neurons, in both mouse visual cortex and macaque V4 [1, 11, 17, 31, 80, 85, 89]. Dynamic modelling of cortical visual responses to natural movies has recently gained attention [12, 82, 87], but temporal predictions of V1 responses and optimised dynamic stimuli remain under-explored. Capturing dynamic processing requires models trained on time-varying inputs such as videos, which introduce additional complexity in terms of model size and data requirements. Recent work has shown that dynamic transformer models are highly effective for video classification tasks [6], and they have begun to be adapted as core models for predicting neuronal activity [5, 82].

Here, we present ViV1T, a movie-trained transformer designed to predict V1 neuronal responses to dynamic visual stimuli. We found that our model outperformed state-of-the-art models in predicting responses to both natural and artificial dynamic stimuli while being faster to run and requiring fewer parameters. Movie-trained ViV1T accurately predicted the well-established selectivity of V1 neurons for orientation and direction, as well as their centre-surround contextual modulation, known to depend on feedback inputs. Beyond its validation on known V1 properties, ViV1T enabled the discovery of new properties. The model predicted that some neurons switched their response to iso-oriented surround stimuli from inhibition to excitation between high and low contrast. By identifying stimuli that maximally drive V1 responses, our model predicted that natural scenes and model-generated stimuli evoked a much broader range of neuronal responses than traditional low-dimensional gratings. Moreover, dynamic surrounds elicited stronger modulatory effects compared to static images. We confirmed these predictions using semi-closed-loop *in vivo* recordings. These results demonstrate that ViV1T is a powerful tool for uncovering the spatial and temporal dynamics of neuronal responses to sensory stimuli.

## 2 Results

### 2.1 ViV1T accurately predicts V1 neuronal responses to dynamic natural stimuli

We trained a transformer-based model, named ViV1T, to predict neuronal responses to dynamic visual stimuli in mouse primary visual cortex (V1). Our model follows the established core-readout framework [44, 45, 48, 87], where a core network receives as inputs the visual stimuli (*e.g*., video) and behavioural information (*e.g*., pupil dilation, pupil centre and running speed), and learns animal-invariant visual and behavioural representations. From these representations, an animal-specific linear readout network [48] predicts calcium responses (Δ*F/F*) of individual neurons. A previous version of the model was developed for predictions of neuronal responses to static images [45]. We extended the model to predict responses to dynamic stimuli. Inspired by the video vision transformer introduced by Arnab *et al*. [6], the core is now made up of two separate transformers: a spatial transformer learns a spatial representation of each video frame, followed by a temporal transformer that captures interactions between frames (see Methods 4.1). Notably, the model does not include any assumption about the connectivity between neurons, neither within V1 nor between V1 and other brain areas.

Here, we compared our model against two published and publicly available state-of-the-art models: the factorised convolutional network (fCNN) [35, 81] and the winning model of the Sensorium 2023 competition DwiseNeuro [10, 82]. We also included a linear-nonlinear model (LN) as a classical baseline [33, 37, 52, 74]. All models were fitted on the Sensorium dataset [81], which consists of calcium recordings of over 78,000 V1 neurons in response to hundreds of 10 s action movie clips from 10 mice. We first evaluated the movie-trained models for their prediction of neuronal responses to new movies (unseen by the models during training). We then tested the models on novel artificial stimuli, including directional pink noise, flashing dots at random spatial positions, drifting Gabor patterns, random moving dots (random dot kinematograms), and flashing static images, all unseen by the models during training [81]. We compared these predictions with the recorded neuronal responses to these stimuli, available for 5 out of the 10 mice (see Supp. Table 4 for the dataset composition). The performance of each model was quantified by using the correlation between predicted and recorded trial-averaged neuronal responses [72] (see Methods 4.3).

Our results showed that ViV1T outperformed the LN model and the two convolution-based models in predicting mouse V1 responses to unseen natural movies, with an average improvement of 26.3% and 5.5% over fCNN and DwiseNeuro, respectively. When testing the generalisation performance to novel artificial stimuli, ViV1T outperformed both LN and fCNN models on all stimulus categories and outperformed DwiseNeuro in 3 out of 5 stimulus categories (see Table 1). Notably, ViV1T showed an average performance drop of 11.2% when behavioural information (running speed and pupil size and position) was withheld from the model, showing the importance of behavioural variables for predicting V1 neuronal responses. We also systematically investigated how much training data and neurons were needed to train ViV1T to reach different levels of performance (Supp. Section S7). As expected, prediction performance improved with more training samples (number of movie clips) and neurons, with number of training samples being the larger limiting factor (Supp. Figure 5).

**Table 1:** Comparison of ViV1T accuracy for V1 neuronal responses to natural and artificial stimuli. Normalised correlation [72] (s.e.m. across animals) between recorded and predicted responses to unseen natural and artificial dynamic stimuli. Avg. shows the normalised correlation averaged across the five stimulus types. The best and second best performing model in each category highlighted in **bold** and underlined, respectively. Supp. Table 1 and Supp. Table 2 show the performance comparison in single trial correlation and trial-averaged correlation.

| Model | Natural movie | Pink noise | Flashing dot | Moving dots | Image | Avg. | Num. Param. |
| --- | --- | --- | --- | --- | --- | --- | --- |
| LN | 0.247<br>(0.013) | 0.173<br>(0.007) | 0.292<br>(0.007) | 0.108<br>(0.006) | 0.282<br>(0.007) | 0.220 | 11M |
| fCNN | 0.551<br>(0.008) | 0.384<br>(0.006) | 0.469<br>(0.006) | 0.394<br>(0.030) | 0.557<br>(0.008) | 0.471 | 11M |
| DwiseNeuro | <u>0.660</u><br>(0.004) | <u>0.556</u><br>(0.006) | **0.648**<br>(**0.009**) | <u>0.568</u><br>(0.019) | <u>0.666</u><br>(0.011) | <u>0.620</u> | 171M |
| ViVIT | **0.696**<br>(**0.009**) | **0.590**<br>(**0.007**) | <u>0.635</u><br>(0.023) | **0.584**<br>(**0.022**) | **0.700**<br>(**0.016**) | **0.641** | 13M |
| ViVIT<br>(without behaviour) | 0.638<br>(0.007) | 0.545<br>(0.004) | 0.511<br>(0.018) | 0.517<br>(0.018) | 0.633<br>(0.011) | 0.569 | 13M |

Importantly, ViV1T also had the best speed *vs*. accuracy performance trade-off (Figure 1C), being more than 20 times faster to run and containing 13 times fewer trainable parameters (13M *vs*. 171M) than the second best performing model (DwiseNeuro); this makes ViV1T more accessible for being used by neurophysiology research groups.

**Figure 1:**
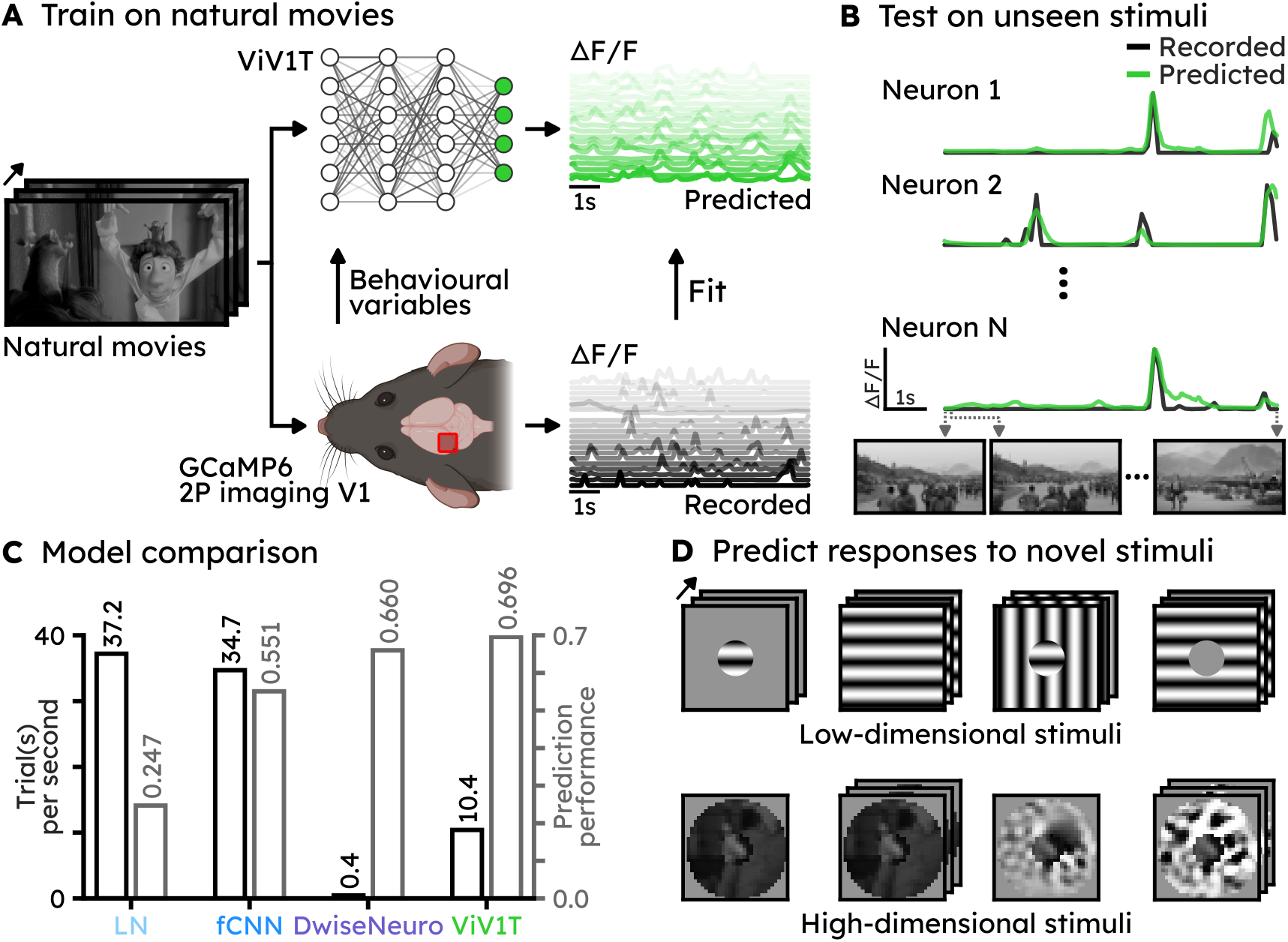
ViV1T accurately predicts V1 neuronal responses to dynamic natural stimuli. (A) Schematic of the predictive model paradigm. ViV1T is trained to predict V1 responses to natural movies. (B) We validate the model by predicting neuronal responses to movies that were not included in the training dataset (test movies). Examples of neuronal responses to a test movie. Predicted responses are in green and *in vivo* recorded responses are in black. Three movie frames are shown below the neuronal traces. (C) Comparison of inference speed (trials per second, in black) and prediction performance (in grey) for V1 responses to natural movies of ViV1T against other state-of-the-art and baseline models. (D) Once validated on traditional stimuli, we use the model to generate dynamic centre-surround most-exciting stimuli, providing novel hypotheses to be tested *in vivo*.

Overall, across the five stimulus categories, ViV1T outperformed current state-of-the-art models (see Table 1), while being more computationally efficient.

### 2.2 Generalisability of ViV1T to new *in vivo* imaging dataset

We tested whether predictions made from the Sensorium dataset were generalisable to other datasets acquired in a different laboratory and including much fewer neurons. To that end, we performed *in vivo* two-photon calcium imaging of mouse V1, imaging neurons during the presentation of natural movies similar to the Sensorium dataset (Methods 4.8). We then tested three different methods to train ViV1T on these new *in vivo* recordings (see Methods 4.8.9). The three methods were: (1) Direct: we initialised a new model (core and readout) and fitted it to the recorded responses to natural movies obtained in our laboratory. (2) Transfer: using the core module trained on the Sensorium dataset, we only initialised a new readout module and optimised it on the new recordings while the core module remained frozen. (3) Fine-tune: similar to the transfer learning setting, but we used a small learning rate (10 times smaller than the original value) to optimise the core module as well. Interestingly, transfer learning achieved prediction performance close to direct training (normalised correlation ± s.e.m.: with transfer learning: 0.468 ± 0.023; direct training: 0.525 ± 0.036) despite the core module in the former setting being trained on recordings that were obtained from a different experimental laboratory (Sensorium dataset). Nevertheless, we found that the fine-tuned models performed the best, with a normalised correlation of 0.611 ± 0.018. Thus, ViV1T generated accurate predictions for our *in vivo* responses even when trained with a limited number of neurons (around 300, see Supp. Table 5) and with the core not fully retrained on our data.

These results align with previous work indicating that pre-training a model on large-scale datasets can improve the model’s performance when fine-tuning it on data obtained in new animals [45, 87]. Altogether, these results show the high generalisability of the model for V1 responses prediction, indicating that the model can be used with limited datasets acquired in different animals and laboratories.

### 2.3 ViV1T accurately predicts orientation and direction tuning curves of V1 neurons

In order to use ViV1T to systematically test which properties of natural stimuli are encoded in V1, we first validated the model by testing whether it accurately predicted the well-established selectivity of V1 neurons for orientation and direction of movement of visual stimuli. We used our movie-trained model to predict responses to drifting gratings and then validated these results by comparing the predicted and the recorded neuronal responses in the Sensorium dataset.

We found that the model accurately predicted single-neuron direction tuning curves in response to drifting gratings (Figures 2A and 2B); both normalised tuning curves and direction selectivity indices were not significantly different between the predicted and recorded neuronal responses (Figure 2C; Wilcoxon signed-rank test: *p* = 0.607). Consistent with this finding, the preferred grating direction of single neurons also closely matched between predicted and recorded responses (Figures 2D and 2E). Notably, these results are not trivially explained by the model being optimised to reproduce single-neuron calcium traces: the model was never trained on responses to gratings and thus learned the orientation and direction tuning properties purely from the neuronal responses to natural movies.

**Figure 2:**
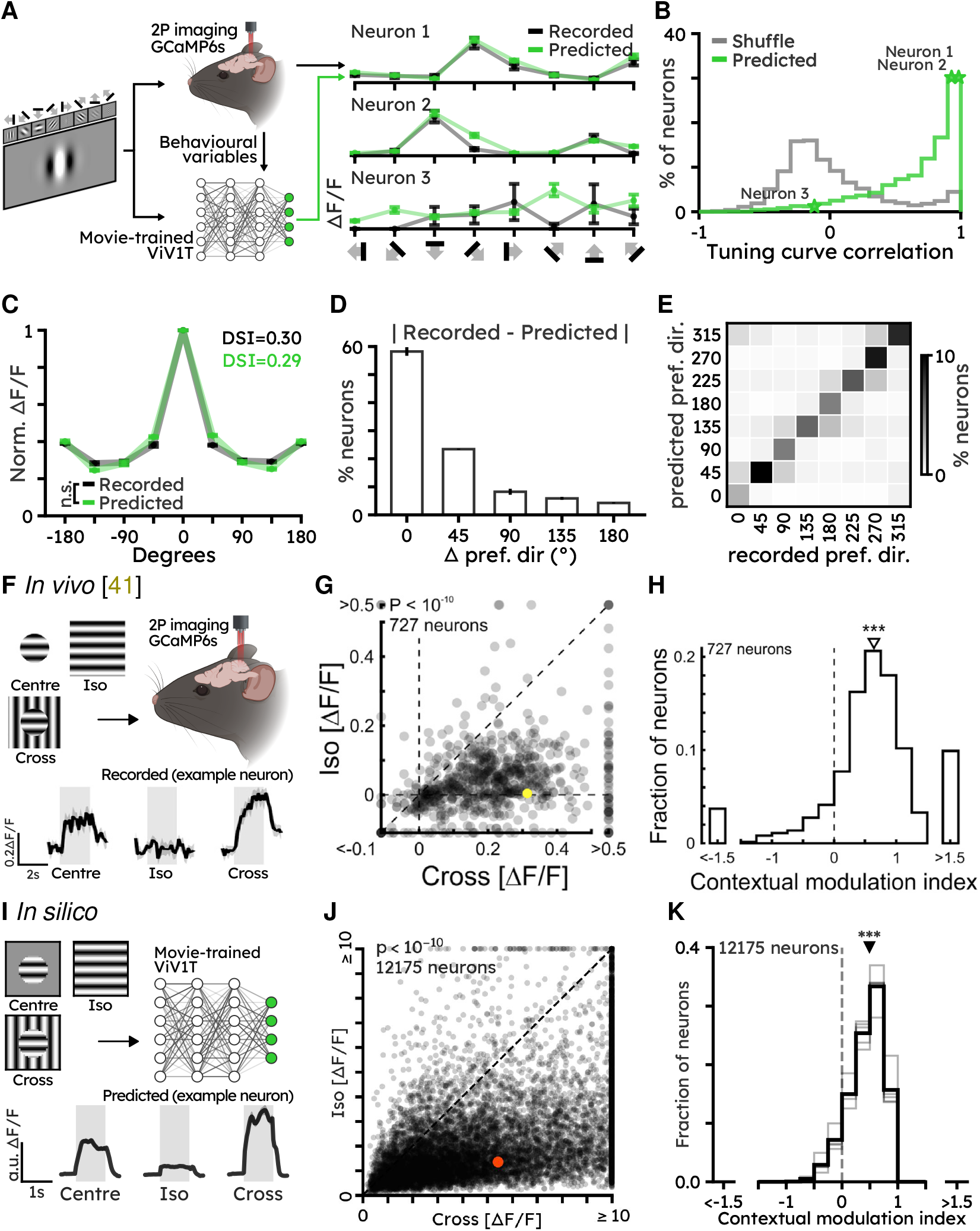
Movie-trained ViV1T predicts orientation and direction tuning and contextual modulation of V1 responses by drifting gratings. (A) Left: Schematic of experimental design. ViV1T was trained exclusively on natural movies and predicted responses to drifting gratings. Predictions were compared to *in vivo* recordings from the Sensorium dataset containing mouse V1 responses to drifting Gabor gratings with eight directions (see Methods 4.2.1). Right: example recorded (black) *vs*. predicted (green) direction tuning curves. (B) Correlation between recorded and predicted direction tuning curves (green line). Grey line indicates results with shuffled recorded tuning curves. (C) Recorded (black) and predicted (green) direction tuning curves, aligned and normalised to preferred direction. Error bars indicate s.e.m. over mice (*N*_neurons_ = 4095 from *N*_mice_ = 3; Wilcoxon signed-rank test n.s.: *p* = 0.607). (D) Angle difference between recorded and predicted preferred direction (*N*_neurons_ = 4095 from *N*_mice_ = 3). (E) Recorded *vs*. predicted preferred direction (*N*_neurons_ = 4095 from *N*_mice_ = 3). (F) Top: Experimental design. V1 neurons expressing GCamP6s were imaged during the presentation of Gabor gratings centred on their receptive field, with grey surround (centre), identical surround (iso), and orthogonal surround (cross) [41]. Bottom: example neuron response for each stimulus (black line is the mean of all trials, shade indicates s.e.m.). (G) Individual neuronal responses to iso *vs*. cross surround gratings (two-sided Wilcoxon signed-rank test *N*_neurons_ = 727 from *N*_mice_ = 9, ***: *p <* 1 × 10^−10^). Red dot indicates example neuron in (F). Reproduced with permission [41]. (H) Contextual Modulation Index (CMI) was computed as the difference divided by the sum of the responses to cross and iso stimuli. Triangles above histograms indicate the median (two-sided Wilcoxon signed-rank test, ***: *p <* 1 × 10^−10^; same neurons as in (G)). All plots from (F – H) reproduced with permission [41]. (I – K) Same as (F – H), but for *in silico* prediction of neuronal responses using Sensorium data (iso *vs*. cross ***: *p <* 1 × 10^−10^; CMI ***: *p <* 1 × 10^−10^).

As a consequence of accurately capturing the orientation tuning curves, ViV1T also reproduced aspects of the spatial organisation of mouse V1 neurons. Recent publications have shown evidence of functional micro-columns in superficial layers of mouse V1 [66, 90] (Supp. Figure 1A and 1B). By quantifying the tuning similarities between neuron pairs as a function of spatial distance (Supp. Figure 1C) [66], we also found evidence for micro-column structure in mouse V1 in the Sensorium dataset: tuning similarity decreased with spatial distance with a substantial drop beyond 50 µm. Applying the same measure to the tuning curves predicted by ViV1T, we found that the model accurately predicts the micro-column structure (Supp. Figure 1C). Notably, the model is not constrained by any assumptions about connectivity between neurons. The spatial organisation is recovered due to the accurate prediction of orientation-selective responses and the known spatial location of the neurons within the imaged region of V1 (from the calcium imaging dataset).

We then compared ViV1T to other models in their ability to predict V1 neurons’ orientation and direction tuning. A particular strength of ViV1T compared to other models is that it accurately captured the full tuning curve shape, and not only the preferred orientation/direction of single neurons (Supp. Figure 2A and 2B). This is shown by comparing the correlations between predicted and recorded direction tuning curves for each of the models (LN, fCNN and DwiseNeuro): our model prediction was the closest to the recorded tuning curves (Supp. Table 3).

Altogether, our results show that ViV1T, exclusively trained on natural movies, accurately predicts V1 neuronal responses to low-dimensional grating stimuli.

### 2.4 ViV1T predicts feedback-dependent centre-surround contextual modulation of V1 neurons

We next explored whether ViV1T could predict contextual modulation of V1 neuronal responses. Numerous previous studies have shown that V1 neuronal responses to a stimulus centred on the neurons’ receptive field can be enhanced or reduced by the presence of a surround stimulus [3, 7, 19, 26, 32, 41, 54, 60, 67, 73, 83, 95]. One characteristic of V1 neurons is that their firing is suppressed when the contextual surround shares the orientation of the centred stimulus: neurons responding to a Gabor patch centred on their receptive field are inhibited when the surround stimulus matches the Gabor’s orientation (iso; Figure 2F). However, these same neurons become more excited when the surround has an orientation orthogonal to the Gabor’s orientation (cross; Figure 2F) [41, 73]. Testing ViV1T with the same stimuli, we found a striking similarity between *in vivo* recordings and the model predictions (Figures 2I to 2K), with stronger responses to cross than to iso stimuli (Figure 2I) [41]. To quantify the effect of this contextual modulation, we used the contextual modulation index (CMI), defined as the difference between iso and cross responses divided by the sum of the responses [41]. ViV1T predicted CMI distributions skewed towards positive values (Figure 2K; mean ± s.e.m. over animals: 0.425 ± 0.025, median: 0.484), similar to what was found *in vivo* (Figure 2H).

Contextual modulation in V1 was previously shown to be partly conveyed through feedback connections from higher visual cortical areas [25, 42], posing the question of whether ViV1T could predict not only feedforward but also feedback responses *in silico*, despite not having any connectivity assumptions in the model architecture. To test this hypothesis, we used visual stimuli (inverse stimuli) that were shown to elicit V1 responses dependent on feedback inputs from higher visual areas. These stimuli consist of large drifting gratings presented with a central patch of uniform mean luminance that varies in size and is centred on the classical receptive field of the neuron. Consistent with previous reports [2, 3, 55], a recent study by Keller *et al*. [42] showed that neuronal responses in V1 demonstrated size-dependent tuning to inverse stimuli that closely resemble responses to classical centred drifting gratings of varying sizes (see Figure 3A). Inverse responses were characterised by significantly longer latencies compared to classical centred gratings responses (Figure 3C) and were markedly attenuated when higher visual areas were suppressed optogenetically [42]. We used ViV1T to predict V1 neuronal responses to the same classical and inverse gratings (Figure 3D). The model was able to predict responses that are strikingly similar to *in vivo* classical and inverse size-tuning curves [42] (Figures 3D to 3F). In addition, ViV1T also predicted the response onset delay to inverse gratings relative to classical gratings (Figure 3F), consistent with the role of feedback inputs in mediating inverse responses [42].

**Figure 3:**
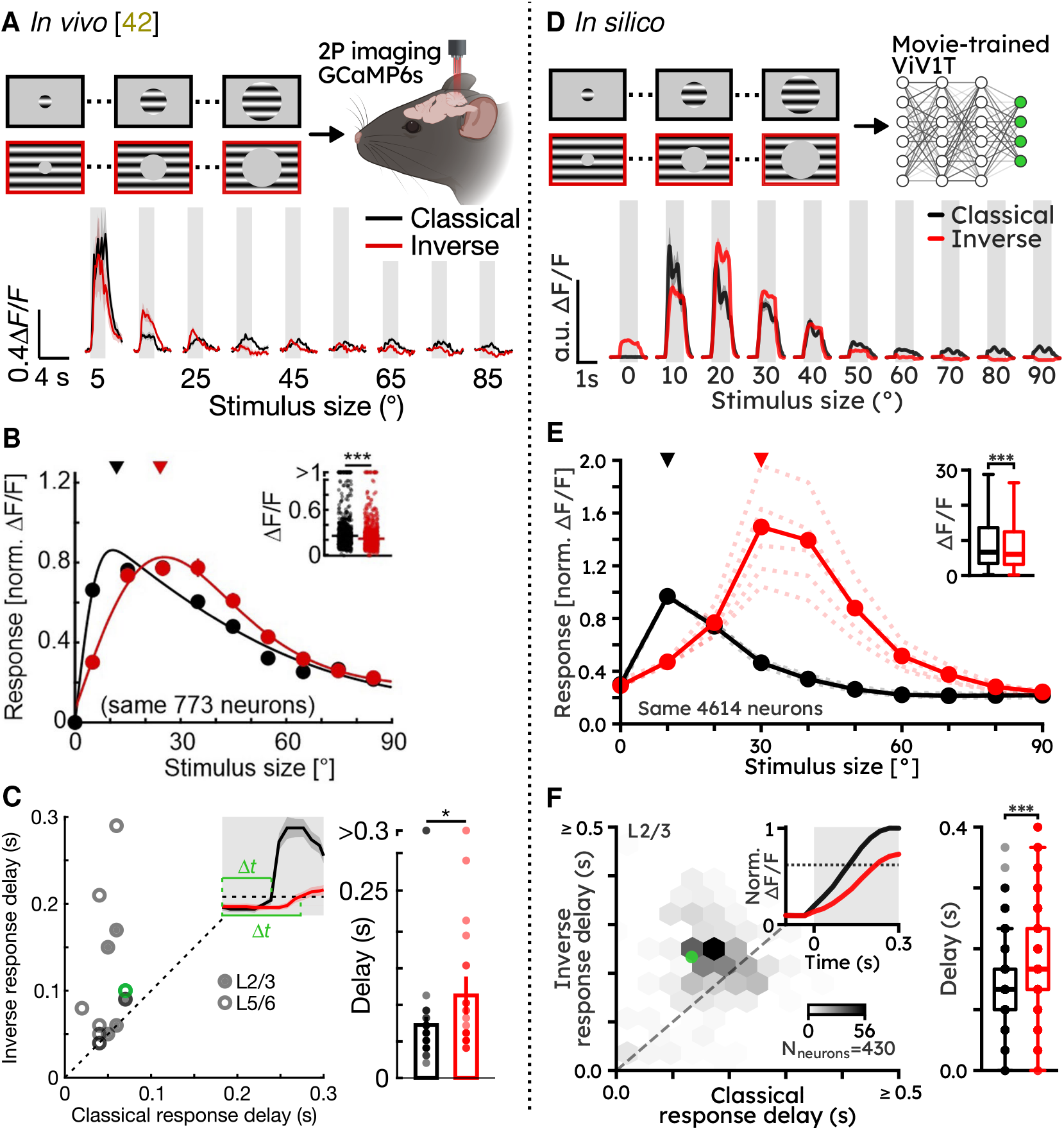
ViV1T predicts feedback-dependent contextual modulation of V1 neurons. Left column: *in vivo* results from Keller *et al*. [42]. All plots reproduced with permission. Right column: *in silico* predictions using Sensorium data. (A) Top: Keller *et al*. [42] imaged mouse V1 neuronal responses to the presentation of classical and inverse Gabor gratings of increasing sizes. Bottom: example trial-averaged responses of a L2/3 neuron for each stimulus size of classical (black) and inverse (red) Gabor gratings. Shaded areas represent stimulus presentation periods. (B) Population-averaged size-tuning curves, normalised to the maximum response to classical stimuli. Solid lines show fits to the data and triangles indicate the median preferred size. The inset shows the maximum responses, with horizontal lines indicating the medians. On average, maximum responses to inverse stimuli were lower than those to classical stimuli (two-sided Wilcoxon signed-rank test: *N*_neurons_ = 773 from *N*_mice_ = 4, ***: *p* = 4.5 × 10^−10^). (C) V1 neurons response delay for classical *vs*. inverse responses. The inset shows example responses to illustrate the difference in response delay to classical and inverse stimuli. The dotted line indicates the response onset threshold (see Methods 4.5.3). Right: Mean response delays of independently tuned neurons (*N*_neurons_ = 51 and *N*_neurons_ = 29 to classical and inverse gratings from *N* _mice_ = 8, two-sided Wilcoxon rank-sum test, *: *p* = 0.031). (D – F) Same as (A – C) but from *in silico* prediction using movie-trained ViV1T; (E) maximum response, two-sided Wilcoxon signed-rank test: *N*_neurons_ = 4614 from *N*_mice_ = 5, ***, *p <* 1 ×10^−10^; (F) average response onset delay, two-sided Wilcoxon rank-sum test: *N*_neurons_ = 1220 and *N*_neurons_ = 852 neurons to classical and inverse gratings from *N*_mice_ = 5, ***: *p <* 1 × 10^−10^.

Comparison with other models showed that all models, other than LN, predicted a contextual modulation index (CMI) distribution skewed toward positive values, consistent with *in vivo* data (Supp. Table 3). All models, except LN, also reproduced size tuning curves for classical and inverse gratings (Supp. Figure 2C and 2D). Notably, only DwiseNeuro and ViV1T learned the onset delays of the feedback receptive fields, with ViV1T being closest to the recorded onset delays (Supp. Figure 2E and Supp. Table 3). Altogether, ViV1T performed either better or as well as other models across the different visual response properties that were tested.

These results show that ViV1T can accurately predict contextual modulation of V1 neuronal responses, which is known to depend on feedback inputs from higher visual areas. Notably, the model predicted the response properties of neurons to surround stimuli solely from their responses to the natural movies used in the training dataset, since there is no built-in mechanism for modelling connectivity between neurons (either within V1 or from feedback inputs) in the model architecture, and no particular bias for inverse stimuli in the training dataset. These results show that ViV1T can capture functional patterns imposed by inter-area feedback connectivity in a purely data-driven fashion.

### 2.5 ViV1T predictions reveal new properties of contextual modulation of V1 neurons

We have demonstrated that movie-trained ViV1T accurately predicts V1 neuronal responses to centre-surround grating stimuli. We then used the model to systematically test which properties of contextual stimuli impact V1 responses and confirmed these predictions with semi-closed-loop *in vivo* calcium imaging.

To that end, we performed *in vivo* two-photon calcium imaging of mouse V1 neurons, with two sessions per field-of-view (FOV) (Methods 4.8). In the first session, V1 neurons were imaged during the presentation of natural movies. We then fine-tuned ViV1T with these neuronal responses and used the model to predict responses to novel stimuli (Methods 4.8.9). Finally, in the second session, responses from the same neurons were imaged in order to verify the *in silico* predictions made by the model.

#### 2.5.1 Low contrast switches contextual responses of a subpopulation of neurons

In natural environments, the contrast of visual inputs naturally varies due to changes in lighting conditions, surface textures, and viewing conditions. How these contrast differences influence contextual modulation in V1 is still not fully understood [7, 19, 25, 28, 38, 68, 69, 73]. We used ViV1T to generate predictions of contextual responses for both high- and low-contrast stimuli. The movie-trained model predicted responses to the same sets of centre-surround grating stimuli, either presented at full contrast or at 5% of the original contrast (Figure 4A).

**Figure 4:**
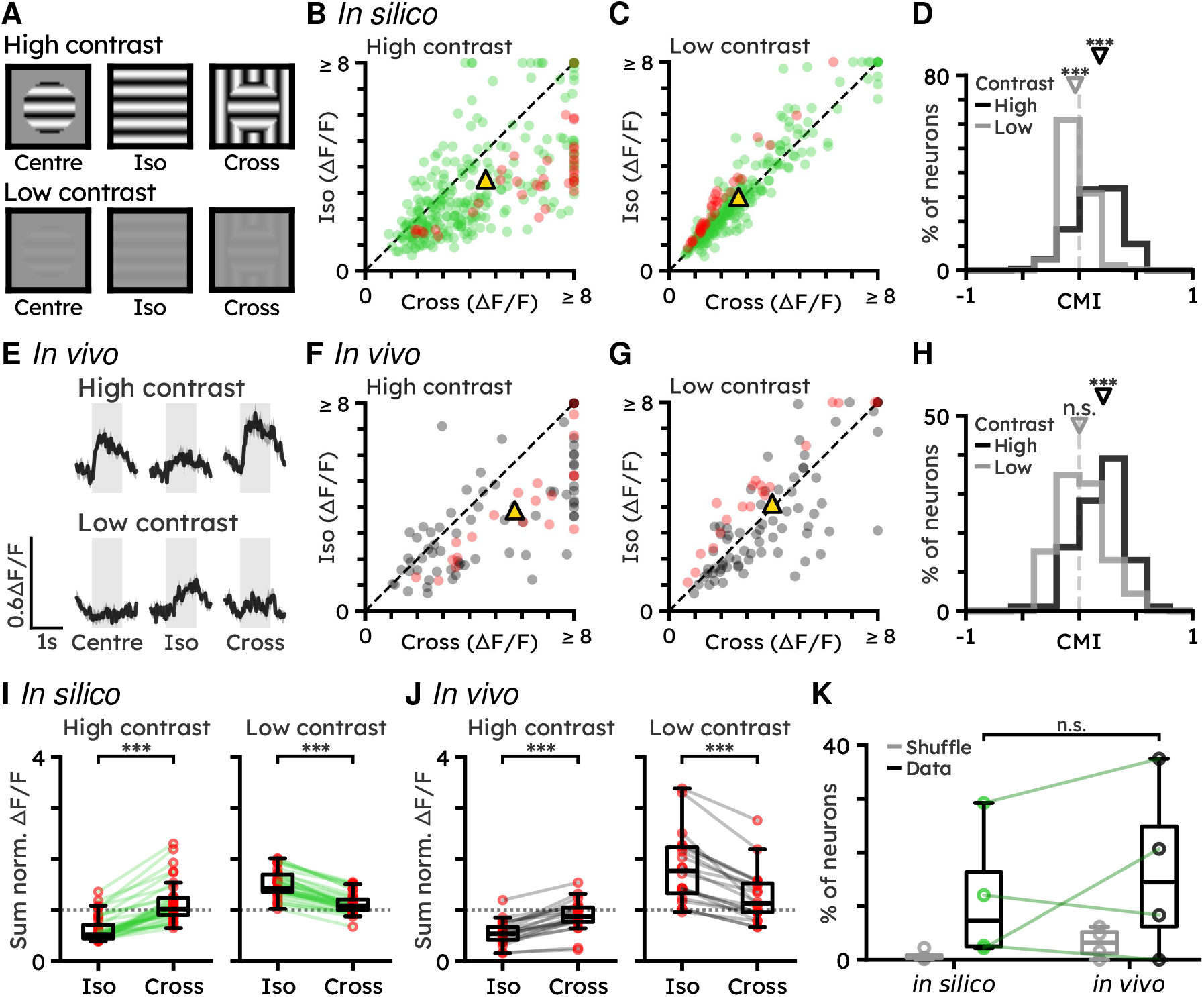
A subpopulation of V1 neurons switches contextual responses with contrast. (A) Illustration of the centre, iso-surround and cross-surround Gabor grating stimuli in high (top row) and low (bottom row) contrast. (B) *In silico* responses to high contrast iso- and cross-surround gratings. Each dot is the trial-averaged Δ*F/F* for each neuron. Gold triangle indicates the population average. Red dots show neurons that switch contextual modulation with contrast (two-sided Wilcoxon signed-rank test: *N*_neurons_ = 303 from *N*_FOVs_ = 4, ***: *p <* 1 × 10^−10^). (C) Same as (B) but with low contrast stimulus (two-sided Wilcoxon signed-rank test: ***:*p* = 8.477 × 10^−8^). (D) Contextual modulation index (high contrast, mean: 0.156, (black triangle) median: 0.181; low contrast, mean: −0.028, (grey triangle) median: −0.036. Two-sided Wilcoxon signed-rank test, high contrast ***: *p <* 1 × 10^−10^, low contrast ***: *p* = 2.342 × 10^−8^). (E) Trial-averaged calcium traces of an example neuron that switches contextual modulation between high and low contrast of iso- and cross-surround grating stimuli. Shaded areas show s.e.m. across stimulus presentations. (G) Same as (B) for *in vivo* recordings (two-sided Wilcoxon signed-rank test: *N*_neurons_ = 92 from *N*_FOVs_ = 4, ***: *p <* 1 × 10^−10^). (F) Same as (F) but with low contrast stimulus (two-sided Wilcoxon signed-rank test ***: *p* = 3.684 × 10^−1^). (H) Same as (D) but for *in vivo* recordings (high contrast mean: 0.189, median: 0.213; low contrast mean: 0.010, median: 0.000. two-sided Wilcoxon signed-rank test, high contrast ***: *p <* 1 × 10^−10^; low contrast n.s.: *p* = 9.472 × 10^−1^). (I) *In silico* neurons that switch contextual modulation between high- and low-contrast stimuli (two-sided Wilcoxon signed-rank test: *in silico N*_neurons_ = 36, high contrast ***: *p <* 1 × 10^−10^, low contrast ***: *p <* 1 × 10^−10^). All responses are normalised by response to centre grating at high contrast (dotted line). (J) Same as (I) but with *in vivo* recordings (two-sided Wilcoxon signed-rank test: *in vivo N*_neurons_ = 20, high contrast ***: *p* = 1.907 × 10^−6^, low contrast ***: *p* = 1.907 × 10^−6^). (K) Percentage of (green) *in silico* and (black) *in vivo* neurons that switch contextual modulation. Each dot indicates the percentage of neurons switching contextual modulation between high and low contrast stimuli per FOV. Grey boxes indicate the results obtained from shuffled responses (two-sided Wilcoxon signed-rank test ***: *p* = 6.250 × 10^−1^).

Consistent with previous studies [25, 68], at lower contrast, the overall effect of contextual modulation was found to be much weaker, with a median CMI of −0.036 at low contrast compared to 0.156 at full contrast (Figure 4D). Interestingly, the model predicted that a small subpopulation of neurons maintained a high CMI but switched their surround suppression towards excitation between high and low contrast settings. About 11.5% ± 6.3% (average ± s.e.m. over FOVs) of the neurons showed a stronger predicted response (≥ 20% Δ*F/F*) to cross-than iso-surround stimuli at high contrast, while they had a stronger predicted response (≥ 20% Δ*F/F*) to iso-than cross-surround stimuli (Figure 4I) at low contrast. Notably, we also found a subpopulation of neurons with similar properties when using the Sensorium-trained model and dataset to predict responses to the same stimuli (Supp. Figure 3).

We validated these predictions with our *in vivo* recordings of V1 neuronal responses. We found that, as predicted and expected, high-contrast stimuli elicited a decreased response for an iso-oriented surround and an increased response for an orthogonally-oriented (cross) surround. The same stimuli presented at low contrast elicited much weaker contextual modulation (Figure 4H). As predicted, we also found that a subpopulation of neurons (average ± s.e.m. over FOVs, 16.6% ± 8.2%) maintained a high contextual modulation but switched their surround inhibition by iso-oriented gratings towards excitation, between high and low contrast stimuli (Figures 4J and 4K). This percentage was not significantly different between the *in silico* predictions and the *in vivo* data, and was higher than the proportion of such neurons from a bootstrap shuffled distribution (Figure 4K).

These results show that ViV1T can be used to identify *in silico* new subpopulations of neurons based on their functional properties.

#### 2.5.2 ViV1T-predicted stimuli enable exploration of the full dynamic range of V1 responses

In addition to predicting neuronal responses to visual stimuli, ViV1T can also be used to generate stimuli optimised for a given neuronal response (*e.g*., maximal response). A key advantage of the *in silico* experiment is that it can be used in a semi-closed-loop [85, 89] fashion to generate and then test high-dimensional optimised stimuli, beyond traditional Gabor gratings and the limited sets of natural images and videos. Methods 4.6 details the procedure for finding and generating most exciting images (MEI) and videos (MEV) for single neurons (*i.e*., the stimulus that will generate the highest average response). Here, we compared how different types of surround stimuli affect neuronal responses to a centre stimulus.

We first used the model to find the grating stimulus eliciting the strongest response for each neuron: the preferred position and drifting direction of a Gabor grating presented during 1 s. We then used the model to test all possible combinations of drifting grating surrounds (8 directions) to find the surround that elicited the strongest increase in response, for each neuron. We repeated the same process to test natural movie surrounds taken from the natural movies used in the first imaging sessions. Finally, we used ViV1T to generate the most-exciting dynamic surround stimuli that would elicit maximal neuronal responses, larger than those evoked by gratings and natural movie surrounds (Methods 4.6). Figure 5A shows the responses of an example neuron to the most-exciting grating centred on its receptive field (grating centre, baseline), as well as the responses to the same centre grating combined with the most-exciting surround grating, natural movie and ViV1T-generated stimuli. Overall, the model predicted that model-generated dynamic surrounds elicited the strongest response, followed by natural movie and grating surrounds, with a 109%, 47% and 33% median response increase over the response to the grating centre, respectively (Figure 5B).

**Figure 5:**
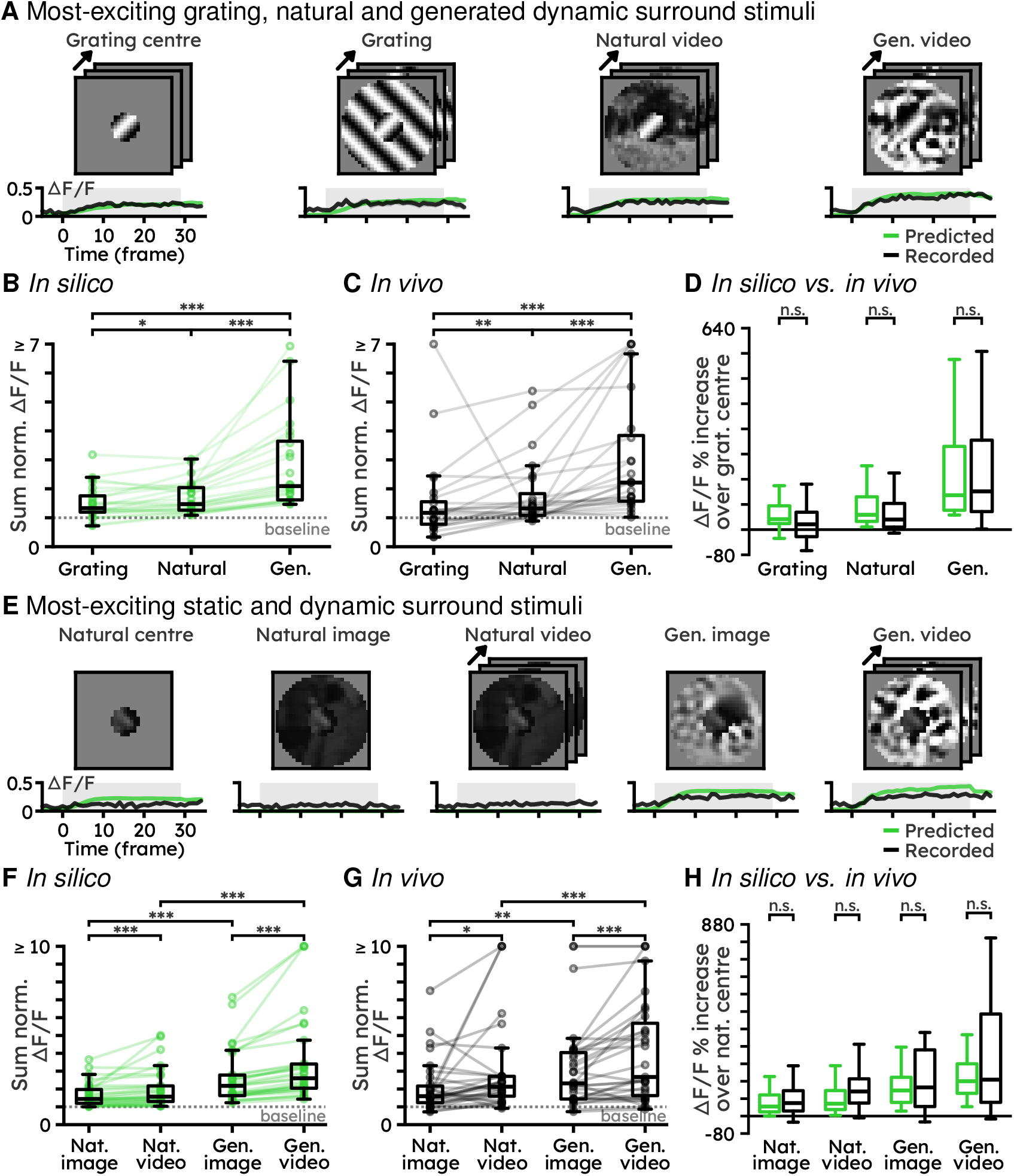
Natural and ViV1T-generated dynamic surrounds elicit stronger contextual modulation than gratings and static surrounds. (A) Example of contextual responses of a single V1 neuron. The left-most column shows the response to the most-exciting grating, centred on the neuron’s receptive field. The right columns show the responses of the same neuron to stimuli combining the same centre grating with the most-exciting grating, natural video and ViV1T-generated video surrounds. The black and green lines are the trial-averaged recorded and predicted Δ*F/F*. (B) *In silico* predicted responses to the different most-exciting dynamic surround. All responses were summed over the stimulus presentation window and normalised to the predicted response to most-exciting grating centre (*i.e*. baseline). Each dot is the trial-averaged response of a single neuron. (One-sided Wilcoxon signed-rank test: *N*_neurons_ = 24 from *N*_FOVs_ = 3. Grating *vs*. natural *: *p* = 1.307 × 10^−2^; natural *vs*. generated ***: *p* = 1.788 × 10^−7^; grating *vs*. generated ***: *p* = 1.788 × 10^−7^). (C) Same as (B) but with *in vivo* recording (Grating *vs*. natural **: *p* = 3.775 × 10^−3^; natural *vs*. generated ***: *p* = 2.503 × 10^−6^; grating *vs*. generated ***: *p* = 1.252 × 10^−5^). (D) Response increase over baseline for *in silico* and *in vivo* responses to different surround stimuli (two-sided Wilcoxon tests between *in silico* and *in vivo* response: grating surround n.s.: *p* = 0.252, natural surround n.s.: *p* = 0.565, generated surround n.s.: *p* = 0.684). (E) Same as (A) but with the most-exciting natural static centre stimulus combined with most-exciting natural static, natural dynamic, ViV1T-generated static and generated dynamic surrounds. (F) *In silico* predicted responses to the different most-exciting dynamic surrounds. All responses were summed over the stimulus presentation window and normalised to the predicted response to most-exciting natural image centre stimuli (*i.e*., baseline). Each dot is the trial-averaged response of a single neuron. (One-sided Wilcoxon signed-rank test: *N*_neurons_ = 31 from *N*_FOVs_ = 3. natural image *vs*. natural video ***: *p* = 3.479 × 10^−5^, generated image *vs*. generated video ***: *p* = 1.863 × 10^−9^, natural image *vs*. generated image ***: *p* = 1.863 × 10^−9^, natural video *vs*. generated video ***: *p* = 1.863 × 10^−9^). (G) Same as (F) but for *in vivo* recordings (natural image *vs*. natural video *: *p* = 1.830 × 10^−2^, generated image *vs*. generated video ***: *p* = 1.556 × 10^−4^; natural image *vs*. generated image ***: *p* = 1.403 × 10^−3^, natural video *vs*. generated video ***: *p* = 1.222 × 10^−5^). (H) Response increase over baseline for *in silico* and *in vivo* responses to different surround stimuli (two-sided Wilcoxon tests between *in silico* and *in vivo* response: natural image surround n.s.: *p* = 1.554, natural video surround n.s.: *p* = 0.317, generated image surround n.s.: *p* = 2.941, generated video surround n.s.: *p* = 1.469).

We validated these predictions with the second session of *in vivo* imaging of V1 responses (see black traces in Figure 5A). Our results show that predictions were accurate: model-generated dynamic surrounds elicited the strongest response, followed by natural movie and grating surrounds, with a 122%, 32% and 17% response increase, respectively (Figure 5C), and no significant difference between *in silico* predictions and *in vivo* recordings (Figure 5D, grating surround *p* = 0.252; natural surround *p* = 0.565; generated surround *p* = 0.684). Additional examples of MEIs and MEVs are available at github.com/bryanlimy/ViV1T-closed-loop.

These results demonstrate that simple stimuli like drifting gratings produce only limited contextual modulation in individual V1 neurons, whereas natural and model-generated surround stimuli evoke much broader response ranges. This suggests that surround orientation represents just one factor among many that influence neuronal contextual responses. ViV1T therefore enables systematic investigation of which high-dimensional natural stimulus properties (and their combinations) drive contextual modulation in V1 neurons.

#### 2.5.3 Dynamic surrounds elicit stronger contextual modulation than static surrounds across stimulus types

We next tested the relative impact of dynamic *vs*. static surrounds on the modulation of V1 neuronal responses. We first used ViV1T to determine for each neuron the preferred static natural centre stimulus (taken from the natural movie training set). We then used the model to predict neuronal responses to stimuli combining this preferred centre stimulus with most-exciting surrounds: (1) natural static and dynamic surrounds (taken from the natural movie training set), (2) ViV1T-generated static and dynamic surrounds (see examples in Figure 5E). The model predicted that dynamic surrounds elicit stronger contextual modulation than static surrounds, with a median response increase of 8% and 24%, for natural and ViV1T-generated stimuli, respectively (Figure 5F).

These predictions were validated by our *in vivo* recordings of V1 responses (Figure 5G). As predicted, both model-generated images and videos evoked responses that were stronger than those evoked by natural images and videos, demonstrating the effectiveness of the method to generate optimised stimuli (natural image *vs*. generated image: *p* = 1.403 × 10^−3^, natural video *vs*. generated video: *p* = 1.222 × 10^−5^). Moreover, both natural and generated dynamic stimuli evoked larger responses than their static counterparts (Figure 5G) (natural image *vs*. natural video: *p* = 1.830 × 10^−2^; generated image *vs*. generated video: *p* = 1.556×10^−4^). There was no significant difference between the predictions and *in vivo* recordings (Figure 5H, natural image surround *p* = 1.554; natural video surround *p* = 0.317; generated image surround *p* = 2.941; generated video surround *p* = 1.469).

We finally tested whether ViV1T could be used to accurately predict not only individual neuron responses but also mean population responses to visual stimuli. We used the model to predict the mean response of all V1 imaged neurons to different visual stimuli. The results were similar to those obtained with single neurons (Supp. Figure 4).

These results show that dynamic surrounds elicit stronger contextual modulation than static surrounds across stimulus types (natural movies and model-generated stimuli), both for single neurons and V1 neuronal populations. ViV1T is thus a powerful tool to investigate response properties not only of single neurons but also of neuronal populations whose average responses can be characterized functionally, for example to determine the output of a subpopulation of neurons to a downstream area.

## 3 Discussion

In this study, we introduce ViV1T, a transformer-based model trained on natural movies to predict neuronal responses in mouse primary visual cortex (V1). Our model outperformed state-of-the-art models in predicting responses to both natural and artificial stimuli, while being significantly more computationally efficient. ViV1T accurately predicted both single-neuron and population responses to both static and dynamic visual stimuli. Although trained exclusively on natural movies, ViV1T accurately captured V1 neuronal tuning properties, including direction tuning, size tuning, and feedback-dependent centre-surround interactions. ViV1T also enabled the discovery of new visual response features, which we confirmed by *in vivo* recordings: (1) a subpopulation of neurons switching their preferred response to surround stimuli when contrast changed from high to low; (2) a wider range of contextual responses elicited by natural and model-generated surround stimuli compared to gratings; (3) dynamic surrounds eliciting stronger contextual modulation than static surrounds. These results highlight ViV1T as a powerful tool for probing spatial and temporal features of neuronal responses to visual stimuli. Notably, ViV1T is fully open-access, with all code, training, and analysis scripts provided, facilitating its use by neurophysiology research groups and ensuring reproducibility across laboratories.

### Temporal predictions of V1 responses: static *vs*. dynamic visual stimuli

This study introduces a novel methodology for generating optimised dynamic stimuli, providing a new framework for systematically probing the temporal properties of V1 neurons, both at the single neuron and neuronal population levels. To the best of our knowledge, the most-exciting dynamic stimuli for V1 responses have not been published before. ViV1T’s ability to synthesise maximally exciting stimuli (most exciting images, MEIs, and videos, MEVs) revealed a broader range of responses not captured by traditional parametric stimuli (gratings), suggesting that ViV1T has learned general principles of visual encoding from the 2 h presentation of videos (action movies). Our results show that one of these principles is that dynamic surrounds elicit stronger contextual modulation than static surrounds across stimulus types (natural movies and model-generated stimuli), highlighting the importance of motion in visual information processing.

Another advantage of temporal predictions of V1 responses is the ability to investigate the interactions between behaviour and visual responses. Our model integrates behavioural signals (locomotion speed, pupil size and pupil centre position), and we show that incorporating these parameters improves prediction performance (Table 1). As such, it is possible to use the model to systematically test the impact of each of these parameters at different timescales. Notably, the model can integrate additional recorded behavioural parameters, such as whisker and snout movements, and head direction.

One potential limitation in exploring the temporal properties of V1 responses in this study, is that we used calcium imaging datasets (Sensorium and our own *in vivo* data). Calcium sensors have a low temporal resolution, preventing predictions in the millisecond temporal domain. With recent advances in modelling neural activities at single-spike resolution using transformers [92, 93], we do not foresee any obstacle in adapting ViV1T to electrophysiological data.

### Using ViV1T to reveal neuronal functional properties

A significant advantage of ViV1T is that it can predict responses to and generate specific stimuli, enabling hypothesis-driven experiments, based on exhaustive stimulus screening *in silico* that would not be possible within the limited time of *in vivo* recordings. Our results show that the range of V1 responses to generated stimuli was significantly higher than those evoked by gratings and by the limited set of natural movies used in the experiments. These results can be used to systematically probe which features of natural surround stimuli drive contextual modulation of V1 neurons. A similar approach could be used not only to find most exciting stimuli, but also to identify other functional characteristics of neuronal populations, for example, by generating stimuli that maximise population entropy or discriminability between neuronal responses.

ViV1T’s ability to predict *in silico* responses to multiple combinations of visual stimuli can also be used to identify new subpopulations of neurons based on their functional properties. We demonstrated this by identifying a subpopulation of neurons that switched their contextual responses to iso-oriented surrounds between high and low contrasts. While requiring *in vivo* validation, these predictions can guide recording strategies, for example, by anticipating the expected proportion of neurons and the characteristics of their responses.

To our knowledge, our study presents the first report of model-generated stimuli optimised for neuronal populations. This approach will be beneficial for predicting and testing population responses (*e.g*., of specific cell-types such as interneurons), and for example, to test different types of outputs of neuronal subpopulations to a downstream area (*e.g*., between V1 and higher visual areas).

### Model architecture and generalisability of ViV1T to novel stimuli and animals

As larger datasets of neuronal activity and computational resources become available, the vision of a foundation model for mouse V1 becomes increasingly realistic [8, 71, 87, 94]. ViV1T provides a scalable architecture that, with continued development, could form the basis for such a model. We suggest that one of the reasons ViViT outperforms state-of-the-art models on visual response predictions and requires fewer parameters is that its self-attention mechanism can better model longrange temporal relationships in spatiotemporal patches than CNN-based models [88]. This approach is thus particularly suitable for modelling temporal responses not only to visual stimuli but also to other sensory stimuli and motor responses.

Our results show that predictions made from the Sensorium dataset were generalisable to our own *in vivo* recordings (*i.e*., performed in a different laboratory). This demonstrates the high generalisability of the model for V1 response predictions and highlights the potential of this approach toward foundation models. In addition, our model framework is not only applicable to visual responses in visual cortex but also to other stimuli, brain areas and species.

Altogether, our results show that data-driven ViV1T is a powerful tool for probing spatial and temporal features of V1 neuronal responses to natural visual stimuli. The open access release of our model, along with the availability of extensive datasets of neuronal responses to sensory stimuli, provides the neurophysiology community with powerful resources to accelerate discoveries in sensory neuroscience.

## 4 Methods

### 4.1 Model architecture

ViV1T follows the recently established approach of the core-readout framework to model visual responses [16, 18, 30, 44, 45, 48, 87]. In this framework, the core module is a large network that learns a joint visual and behavioural presentation across animals, and a small per-animal readout module predicts individual neuronal activities from the core presentation. In particular, ViV1T is composed of a transformer-based core module followed by animal-specific linear readout modules. Figure 6 illustrates the architecture of the model.

**Figure 6:**
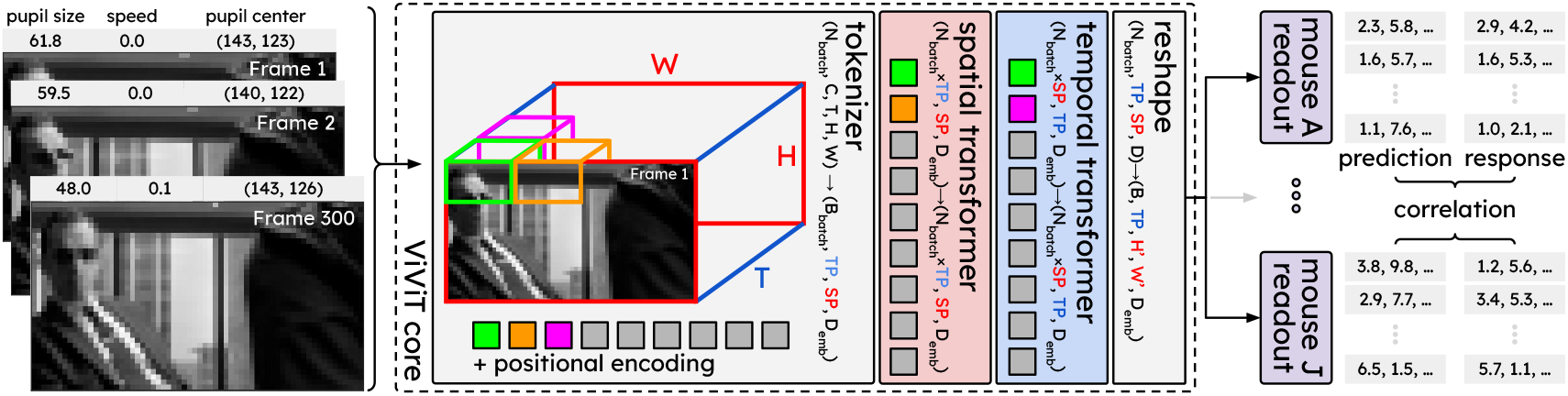
ViV1T architecture. The core network receives the video and behaviour information (pupil size, speed, pupil centre) of each trial and learns a joint spatiotemporal latent representation across all animals via two separate transformers over the spatial and temporal patches of the stimulus. The per-animal readout network then receives the latent representation as input and predicts the response of each recorded neuron. We compute the single trial correlation between the predicted and recorded response to evaluate the model.

#### 4.1.1 ViV1T core module

Based on the video vision transformer developed by Arnab *et al*. [6], the ViV1T core module consists of three main components: (1) a tokeniser, (2) a spatial transformer and (3) a temporal transformer.

##### Tokeniser

The tokeniser extracts spatiotemporal tubelet patches (or tokens) of the input video and behavioural variables (*i.e*., horizontal and vertical coordinates of the pupil centre, pupil dilation and running speed) and learns an embedding of each patch. To combine 4-dimensional visual (*i.e*. `(T=300, C=1, H=36, W=64)`) and 2D behavioural (*i.e*. `(T=300, B=4)`) inputs, we broadcast each behavioural variable to match the height and width of the video (*i.e*. `(T, B, H, W)`). We then concatenate the broadcasted behavioural variables with the video in the channel dimension (*i.e*. `(T, C + B, H, W)`). The tokeniser then uses a sliding window to extract 3-dimensional tokens over spatial and temporal dimensions jointly (*e.g*. light green, orange and pink boxes in Figure 6), where the spatial and temporal patch sizes (*P*_spatial_ and *P*_temporal_) and strides (*S*_spatial_ and *S*_temporal_) are hyperparameters of the model. This results in a 8-dimensional vector `(C + B, TP, SP = SP`_*H*_ × `SP`_*W*_, *P*_temporal_, *P*_spatial_, *P*_spatial_), where TP and SP are the number of temporal and spatial tokens extracted. We then flatten the 3-dimensional tokens and channels into `(TP, SP,` *P*_temporal_ × *P*_spatial_ × *P*_spatial_× (*C* + *B*)). We zero-pad the input video to ensure that all frames are covered, regardless of the spatial and temporal patch sizes and strides. We learn the token embeddings by applying layer normalisation and a fully connected layer with hidden dimension *D*_emb_, resulting in an embedding of shape (TP, SP, *D*_emb_) for a single sample. Finally, we employ patch dropout [75], which randomly zeros out some of the token embeddings with a dropout rate of *p*_token_, to avoid overfitting.

##### Spatial transformer

The spatial transformer operates over the spatial patches (*i.e*. green and orange patches in Figure 6) to learn the spatial relationships in each frame. To apply the self-attention mechanism only to spatial patches, we combine the batch dimension with the temporal token dimension, *i.e*. from (*N*_batch_, TP, SP, *D*_emb_) to (*N*_batch_ × TP, SP, *D*_emb_). The transformer itself consists of multiple attention blocks [84], where the number of these blocks is a hyperparameter of the model. We also integrated several recently proposed improvements to the transformer architecture into our transformer core. We replaced the standard self-attention block with parallel attention [86], which fuses the `QKV` projection layer with the feedforward module and thus reduces the number of operations without sacrificing performance [24]. In addition, we used the highly hardware-optimised FlashAttention-2 [21] implementation of the scaled-dot-product attention layer to speed up model training and inference. Darcet *et al*. [22] observed that vision transformers often learn redundant artefacts, which can cause some low-information patches to have abnormally high attention values. To improve interpretability of the attention maps, we tested `register` tokens [22]. These learnable tokens (or patches) are appended to each attention layer and prevent the model from allocating high attention scores to random tokens. Finally, we tested various positional encoding methods, including learnable positional encoding, sinusoidal positional encoding, and rotary position embeddings (RoPE) [78, 79], which we then selected as another hyperparameter.

##### Temporal transformer

The temporal transformer receives the embeddings from the spatial transformer as input, where we rearrange the dimension to (*N*_batch_ × SP, TP, *D*_emb_), so that the selfattention mechanism is being applied over the temporal dimension. The temporal transformer has the same architecture as the spatial transformer, with two exceptions: (1) the number of attention blocks is a separate hyperparameter from the spatial transformer, (2) we accounted for the fact that taking into account information from future frames is not physiologically plausible. Since ViV1T predicts the entire response to video input at once, instead of autoregressively, we use a causal attention mask in the temporal transformer to mask out future patches: to predict response *y*_*t*_ at time *t*, the selfattention layer only has access to patches from `TP`_*τ*=0_ to `TP`_*τ*=*t*_ while future patches `TP`_*τ≥t*+1_ are set to zero. The output of the temporal transformer is rearranged back to a 4-dimensional shape (*N*_batch_, TP, SP, *D*_emb_) in order to match the Gaussian readout input dimension (see Methods 4.1.2). Unlike the original video vision transformer, ViV1T does not reduce the dimensionality of the data using the cls token in each transformer.

We conducted an exhaustive Bayesian hyperparameter optimisation to find the best configuration. More details are provided in Methods 4.2.2.

#### 4.1.2 ViV1T readout module

Our readout module follows the method described in Lurz *et al*. [48]. To compute the neuronal response of neuron *i* from mouse *m* with *N*_*m*_ neurons, the module 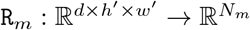 computes a linear regression of the core representation ***z*** with weights 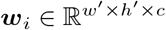, followed by an ELU activation with an offset of 1 (*i.e*. ***o*** = ELU(`R`_*m*_(***z***)) + 1), which keeps the response positive. The regression is performed by a Gaussian readout, which learns the parameters of a 2d Gaussian distribution whose mean ***µ***_*i*_ represents the centre of the receptive field of neuron *i* in the image space and whose variance quantifies the uncertainty of the receptive field position. This uncertainty decreases with training. The response is thus obtained as a linear combination of the feature vector of the core at a single spatial position, which allows the model to greatly reduce the number of parameters per neuron in the readout. Notably, to learn the position ***µ***_*i*_, the readout couples the recorded cortical 2d coordinates of each neuron with the estimated centre of the receptive field from the readout. Moreover, a shifter module is introduced to adjust (or shift) the ***µ***_*i*_ receptive field centre of neuron *i* to account for the trial-to-trial variability due to eye movement [30]. The shifter network ℝ^2^ → ℝ^2^ consists of 3 dense layers with hidden size of 5 and tanh activation; it takes as input the 2d pupil centre coordinates and learns the vertical and horizontal adjustments needed to shift ***µ***_*i*_. We have also tested other previously published readout architectures as part of the hyperparameter optimisation, including factorised and attention readouts [44, 62], but without observing improvement in prediction performance.

### 4.2 Model training and hyperparameter optimisation

#### 4.2.1 Sensorium 2023 dataset

For initial model training and assessment, we used the publicly available Sensorium 2023 dataset [81]. The dataset contains two-photon imaging data from excitatory neurons in superficial layers (200 µm to 425 µm) of the primary visual cortex in awake, head-fixed, behaving mice, using calcium indicator GCaMP6s. Changes of fluorescence over time were extracted for each neuronal soma [81] and resampled from 8 Hz to 30 Hz to match the frame rate of the visual stimuli. In addition, the dataset contains measurements of four behavioural variables: locomotion speed, which is recorded from a cylindrical treadmill at 100 Hz and resampled to 30 Hz, as well as pupil size and horizontal and vertical pupil centre position, which are extracted from tracked eye camera video at 20 Hz and also resampled to 30 Hz. In total, the Sensorium dataset comprises ten imaged field-of-views (FOVs) in ten animals, including the responses of 78,853 neurons to a total of around 1200 min presentation of dynamic visual stimuli (around 120 min of visual stimulation per mouse).

While all mice were imaged during the presentation of natural movies, different mice were shown different artificial stimuli. The calcium responses to the test stimuli are publicly available for only 5 out of 10 mice (mouse `A` to `E,` see Supp. Table 4 for details about the test set composition for each mouse). Each natural movie presentation included 360 movies shown one time and 18 movies shown 10 times, in both cases drawn equally from the “cinematic” and “Sports-1M” classes from the library described in [20]. Three mice (`B, C,` and `E`) were also imaged during the presentation of drifting gratings (Gabors, 8 directions, 3 spatial frequencies, 3 temporal frequencies, Supp. Table 4), with each stimulus presented for 833 ms (25 frames) with ten repetitions [81].

The dataset is already split into training, validation and test sets by the Sensorium 2023 competition organisers. We used the same splits to train, validate and test the models presented in this work.

#### 4.2.2 Model training

The model was trained from scratch on the training data from all 10 mice in the Sensorium dataset. The video inputs, behavioural variables and recorded responses are standardised using the mean and standard deviation measured from the training set. The shared core and per-animal readouts are trained using the schedule-free AdamW optimiser [23] to minimise the Poisson negative log-likelihood between the recorded *y*_*i,t*_ and predicted 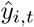 responses over *N* trials and *T* = 300 time points:

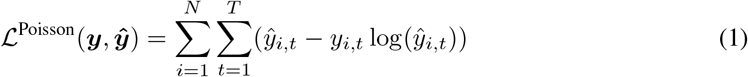

A small value *ε* = 1 × 10^−8^ was added to both ***y*** and ***ŷ*** prior to the loss calculation to improve numeric stability. In line with previous work [45], we found that accumulating the gradients from all mice before a single gradient update step significantly improves model performance as compared to updating the core and one readout at a time. The single trial correlation between *y*_*i*,:_ and *ŷ*_*i*,:_ in the validation set is computed after each training epoch. We stopped the training if the model did not improve over 10 consecutive training epochs. In most cases, the model stopped improving after 100 training epochs. To prevent overfitting, we added dropout [75] in the tokeniser (patch-wise dropout), as well as in each self-attention and feedforward layer in the transformer blocks. In addition, stochastic depths [36] and weight decay were employed. To find the optimal settings for the model over this vast hyperparameter space, we conducted a large Hyperband Bayesian optimization [46] to find the hyperparameters that achieved the best single-trial correlation on the validation set. The initial search space and final hyperparameter settings are detailed in Supp. Table 6. Moreover, the importance of each hyperparameter and the correlation between each hyperparameter and the optimisation objective are provided in Supp. Table 7. The resulting model contains about 13 million trainable parameters.

The LN and fCNN models were trained under the same conditions. The LN model has the same architecture as the fCNN model except that only the output layer includes a non-linearity. Furthermore, ViV1T and fCNN only differ in the core architecture, while the shifter network and readout networks are the same. DwiseNeuro was originally trained with a 7-fold cross-validation on all available data, including the test sets [10, 82]. For a fair comparison and to avoid overfitting, we modified the original DwiseNeuro model training code (github.com/lRomul/sensorium) to fit the model only on the training set. All models were trained from scratch on a single NVIDIA H100 80GB GPU, and all subsequent analyses were performed on a single NVIDIA RTX 4070 Ti SUPER 16GB GPU. The model training and evaluation pipeline is implemented in Python 3.12 and PyTorch 2.8 [61].

#### 4.2.3 Model predictions to novel stimuli (without behavioural variables)

When predicting responses to novel visual stimuli that were not part of the Sensorium dataset, behavioural information (*i.e*. pupil centre, pupil dilation and running speed) was not available. However, most models were trained with and require behavioural inputs to make predictions. To address this issue, we used the average behavioural variables recorded in the training set, *i.e*. we average running speed over the entire training set to get a single value and broadcast this value to match the duration of the novel visual stimulus.

### 4.3 Model evaluation

We evaluated the predictive performance of the models using the normalised correlation *r*_norm_ as proposed by Schoppe *et al*. [72]:

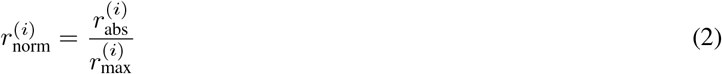

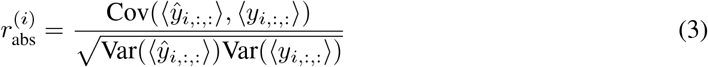

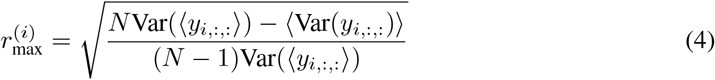

where *y*_*i,j,t*_ and *ŷ*_*i,j,t*_ denote recorded and predicted responses, respectively, for neuron *i* on trial *j* at time *t*, Cov(·, ·) and Var(·) were calculated across trials/stimulus repeats and ⟨·⟩ denotes average across time. *r*_abs_ is the Pearson correlation coefficient between average *in silico* and *in vivo* responses. *r*_max_ is the upper bound of *r*_abs_ achievable by any predictor given the *in vivo* variability of the neuron and the number of trials *N*. *r*_norm_ was calculated independently per neuron and then averaged across all neurons to obtain a population metric.

### 4.4 Analysis of orientation and direction tuning curves

#### Tuning curves

For each neuron, we computed orientation and direction tuning curves by averaging the neuron’s responses to grating stimuli with given orientations/directions across all spatial and temporal frequencies and across trials.

#### Selectivity indices

We computed orientation and direction selectivity indices (OSI and DSI) as follows [87]: 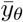 denotes a neuron’s mean response to angle *θ*, averaged over repetitions of that angle,

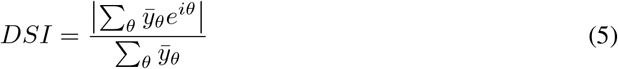

DSI were computed for recorded and predicted tuning curves separately.

For all orientation and direction tuning analyses, we only included neurons that had a higher DSI than the one obtained from shuffled responses. For this, for each neuron, we permuted the averaged responses to each grating 100 times. We computed the DSIs for each of the resulting shuffled responses and used the Gaussian 90-percentile as the threshold.

#### Tuning similarity

We computed the tuning similarity between two neurons as the Pearson correlation between their orientation or direction tuning curves [66]. To compare the similarity of tuning across planes and cortical distance (Figure 2), we grouped neurons from imaged planes separated by 25 µm, to obtain large enough sample sizes. We considered planes at depths 0 µm to 25 µm as the first plane group, 50 µm to 75 µm as the second plane group, and so on to obtain five joined planes altogether from the 10 imaged planes of the Sensorium dataset.

#### Tuning similarity decay

To compare the recorded *vs*. predicted decay in tuning similarity as a function of cortical distance, we fitted decaying exponentials, parametrised by variables *a, b* and *c*, to the tuning similarity curves as a function of cortical distance *d*, but averaged over plane distances Δ. Assuming additive Gaussian noise, we estimated the variance using the maximum likelihood estimate:

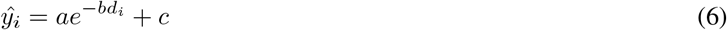

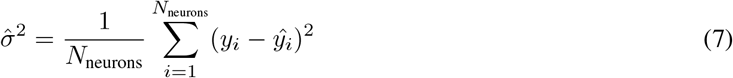

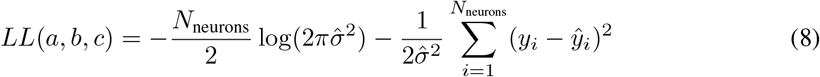

We fitted this model separately to the recorded data, to the predicted data, and both combined. We then computed the log-likelihood ratio, which (under the null hypothesis) follows a *χ*^2^ distribution, to test whether the separate fits significantly improve over the combined fit.

### 4.5 Analysis of centre-surround contextual modulation responses

#### 4.5.1 Estimating artificial receptive fields

To get an estimate of a neuron’s receptive field, we presented the trained model with 100k static fullfield white noise images. Each image was presented for 1 s (*i.e*. 30 frames). To match the length of trials in the Sensorium dataset of 300 frames, we added a grey screen before and after each image presentation: 135 frames of grey screen, followed by 30 frames of white noise and followed by another 135 frames of grey screen to construct a single trial.

The artificial receptive field (aRF) of a neuron *i* is the weighted average of all white noise images, weighted by the summed responses of the neuron to the corresponding image during the presentation window, consisting of *T* = 30 time points:

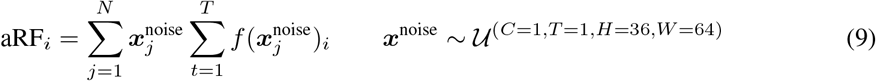

where *f* the model, 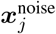 is the *j*^th^ white noise image, *N* is the total number of white noise images, and *f* (***x***_*j*_)_*i*_ is the corresponding response of neuron *i* to 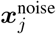. We then standardised the aRF and took its absolute values (*i.e. aRF* = |(*aRF* − *µ*)*/σ*| where *µ* and *σ* are the mean and standard deviation of the values of the aRF). Finally, we fitted a 2D Gaussian to estimate an idealised shape of the aRF, which can be characterised by the centre (*i.e*. the 2D mean of the Gaussian) and the shape (*i.e*. the 2 × 2 covariance matrix of the Gaussian) of the aRF. We repeated this process for each predicted neuron. Since not all aRFs had a good Gaussian fit, we excluded the 5% neurons with the largest aRFs as quantified by the maximum variance of their covariance matrix. Figure 7A shows the aRFs of randomly selected neurons from `mouse` A, where the red circles illustrate the Gaussian fits. Figure 7B shows the aRF centres of all neurons from `mouse` A, which align with our expectation that the population receptive field is at the centre of the monitor, as described in [81]. We used the centre of the 2D Gaussian fit result as the preferred position of the neuron in all subsequent analyses.

**Figure 7:**
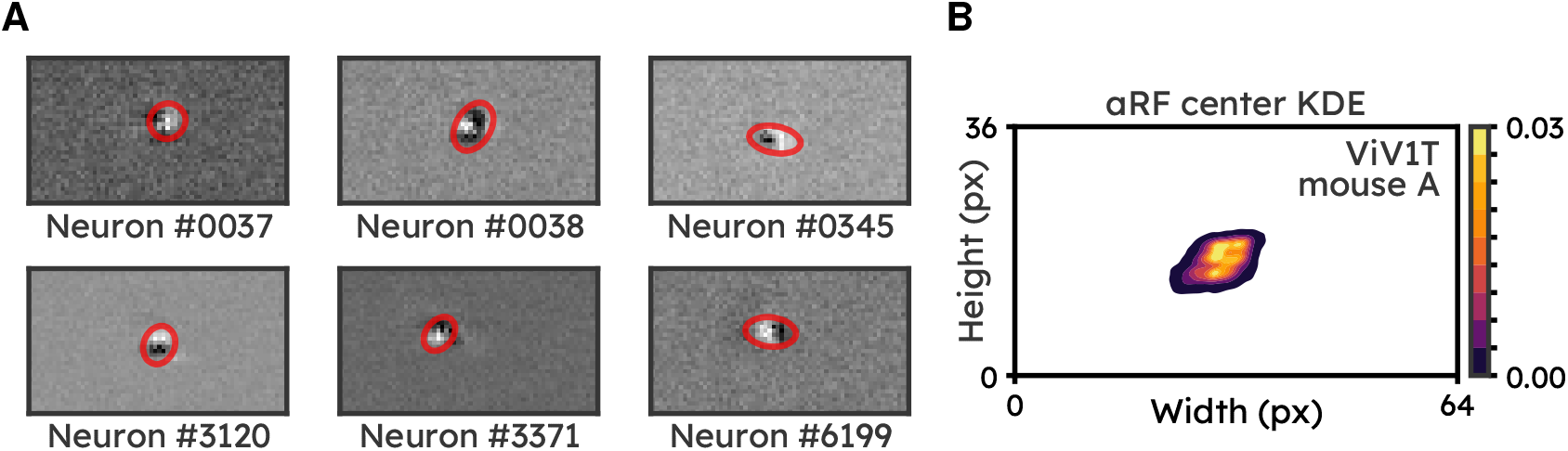
Estimating artificial receptive fields. (A) Estimated artificial receptive fields (aRFs) of six randomly selected neurons from mouse `A.` Red circles illustrate the 1 standard deviation ellipse of the 2D Gaussian fit of each aRF. (B) KDE of the fitted Gaussian centres across the 95% best fitting neurons in `mouse A.`

#### 4.5.2 Centre-surround contextual modulation with drifting gratings

We used the same grating stimuli as described in Keller *et al*. [41]: (1) centre: horizontal grating of size 20^*°*^, spatial frequency of 0.04 cycles per degree and temporal frequency 2 Hz; (2) iso-surround stimulus: centre and surround gratings of the same orientation and direction; (3) cross-surround stimulus: centre and surround gratings of orthogonal orientation. We positioned the centre of each stimulus at the centre of the aRF (Methods 4.5.1) of each neuron. Each stimulus was presented for 1 s (*i.e*. 30 frames), and preceded and followed by 0.5 s of grey screen between presentations. All stimuli were presented at least 12 times in random order. In the low contrast experiments, we used the same stimuli but at 5 % of the original contrast.

In the analysis, we included L2/3 neurons with a preferred stimulus size within 10° of the presented stimulus. We then quantified contextual modulation using the contextual modulation index [41], defined as the difference between the activity to cross and iso stimuli divided by the sum of the two.

#### 4.5.3 Feedforward and feedback receptive fields

##### Size tuning curves

For each neuron, we generated patches of drifting Gabor gratings centred on the neuron’s receptive field (classical stimuli). In addition, we generated inverse stimuli consisting of large drifting gratings presented with a central patch of uniform mean luminance that varies in size and is centred on the classical receptive field of the neuron [41]. As described in Keller *et al*.’s [41], we used a temporal frequency of 2 Hz, spatial frequency of 0.04 cycles per degree, 100 % contrast, and only horizontal gratings moving either up or down. The initial phase of the gratings was randomised. We increased the patch size from 0° to 90° in intervals of 10°. Each of the 40 unique stimuli (up or down, classical or inverse, and 10 possible sizes) was shown for 1 s (30 frames), and preceded and followed by 0.5 s of grey screen between presentations, presented 12 times and in randomised order. For each neuron and per patch size, we computed the average response over the 24 repetitions (across both drifting directions). We only considered L2/3 neurons, and we excluded neurons from further analysis if they did not significantly respond to at least one classical stimulus of any size (z-score *>* 1.96, *p <* 0.05).

##### Response onsets

Following Keller *et al*. [42], we showed classical and inverse gratings to the model. Each stimulus was presented for 0.5 s with 1,000 repeats and randomised initial phases. Response onset to a classical or inverse stimulus is defined as the first time point after stimulus onset where the response passes a z-score threshold of 1.0. We excluded neurons whose response did not exceed the response threshold and whose preferred stimulus size was out of the presented stimulus size of 20° (*i.e*., either smaller than 10° or larger than 30°).

### 4.6 Most-exciting stimuli for single neurons

In this section, we describe the procedure to extract or generate the stimuli that maximally evoke *in silico* responses of a single neuron, which were then verified *in vivo*, as shown in Results 2.5.2. All *in silico* stimuli were presented for 1 s (30 frames), and preceded and followed by 0.5 s of grey screen between presentations. All stimuli were restricted to have a maximum coverage of 60° as we only focus on near-surround contextual modulation [56]. The predicted response of a neuron to a stimulus was computed as the summed response (Δ*F/F*) over the presentation window and averaged over trials.

#### Finding the most-exciting grating centre (dynamic)

We presented drifting grating centres to movie-trained ViV1T to select the most-exciting direction for each neuron of interest. All sinusoidal grating patterns were presented at full contrast, with a spatial frequency of 0.04 cycles per degree, a temporal frequency of 2 Hz, and randomly initialised phases. The grating patch had a stimulus size of 20° and was adjusted to the centre of the neuron’s aRF (see Methods 4.5.1). In total, 8 grating directions were presented in random order, each with at least 10 repeats. We selected the most-exciting grating centre as the pattern that evoked the largest summed response over the stimulus presentations.

#### Finding the most-exciting natural image centre (static)

We extracted image frames from the natural movies in the training data (see Methods 4.8.6) using a sliding window over the temporal (frame) dimension, with a step size of 10 frames. To find the most-exciting natural static centre, we centre-cropped the image to a coverage of 20° and positioned it at the centre of the neuron’s aRF. More than 11,000 movie frames extracted from the 410 movies from the training set were presented to each neuron. Similarly to finding the most-exciting grating centre, we selected the most-exciting natural image centre as the image patch that evoked the largest summed response over the stimulus presentation.

#### Finding the most-exciting grating surround (dynamic)

To find the most-exciting grating surround pattern, we presented different sinusoidal grating surrounds to the model, while fixing the preselected most-exciting grating centre. The surround gratings have a coverage size of 60°, while using the same contrast, spatial and temporal frequencies as the most-exciting grating centre. We selected the surround grating pattern that elicited the largest summed response over the stimulus presentation.

#### Finding the most-exciting surround natural image (static)

As described for the most-exciting natural image centre (static), we presented the movie frames extracted from the training data using a sliding window, with a step size of 10 frames. The movie frames were cropped to a coverage size of 60° with the centre of the image replaced by the pre-selected most-exciting centre pattern. We selected the surround natural image pattern that elicited the largest summed response over the stimulus presentation.

#### Find the most-exciting surround natural video (dynamic)

Similar to finding the most-exciting surround natural image, we presented natural surround dynamic stimuli extracted from the training data with the centre of the stimuli fixed to the pre-selected most-exciting centre. However, instead of extracting movie frames, we extracted 1 s movie clips from the training movies using a sliding window with a step size of 10 frames. About 11,000 1 s movie clips extracted from 410 training samples were presented to each neuron. We selected the surround natural video pattern that elicited the largest summed response over the stimulus presentation.

#### Generating the most-exciting surround image (static)

To generate the most-exciting image (MEI) surround that elicits maximal neuronal responses, we adapted the feature visualisation libraries Lucid [57] and Lucent (github.com/greentfrapp/lucent). As the search space of stimuli is vast, we can assist the stimuli optimisation process by selecting a starting point from the natural images database. To this end, we initialised the image ***x***^MEI^ ~ 𝒰^(*C*=1,*T* =1,*H*=36,*W* =64)^ with the most-exciting natural image that was extracted previously, for each neuron. To ensure that each pixel of ***x***^MEI^ was in the range [0, 255), we applied the sigmoid function element-wise to each pixel prior to presenting the image to the model. Moreover, we applied a circular mask of size 60° to ***x***^MEI^ to replace the far-surround pixels with grey colour. We then predicted the response of neuron *i* using the model ***ŷ***^MEI^ = `ViV1T`_*i*_(255 × `sigmoid` (***x***^MEI^)). To maximise the neuron’s response to the stimulus, we minimised the negative predicted response summed over the presentation window *T* = 30 (1 s):

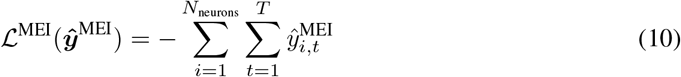

We used AdamW [47] to optimise ***x***^MEI^ using a learning rate of 5×10^−2^ for 200 steps. Using gradient descent to optimise pixel values is prone to high-frequency noise [57]. To alleviate this issue, we initialised and optimised the image ***x***^MEI^ in the frequency domain instead of the spatial domain. Before applying the sigmoid activation and presenting the image to the model, we applied a low-pass filter to cut out half of the spatial frequencies in the image. Then we applied the inverse Fourier transform to convert the ***x***^MEI^ to pixel space. The learning rate, number of optimisation steps and spatial frequency cut-off rates are hyperparameters selected via a grid search and visual inspection of the resulting MEI. A MEI took about 2 min to generate on a single NVIDIA RTX 4070 TI SUPER GPU.

#### Generating the most-exciting surround video (dynamic)

Similar to generating the most-exciting image surround, we also generated a video that maximises the single neuron response over the stimulus presentation window of 1 s. However, instead of creating a static image that was presented for a fixed duration, we initialised the most-exciting video (MEV) ***x***^MEV^ ~ 𝒰^(*C*=1,*T* =30,*H*=36,*W* =64)^ with the most-exciting natural video surround that was extracted previously, for each neuron. We then optimised ***x***^MEV^ using the same objective as in Equation (10) under the same training procedure. Moreover, we only applied the low-pass filter over the spatial dimension of the video, but not the temporal dimension, to allow for more variation in the temporal domain. Similar to generating MEIs, a MEV took about 2 min to be generated.

#### Selecting neurons with high response reliability

As we could only test a limited number of stimuli during the *in vivo* imaging sessions, we selected the top 25 most reliable neurons to generate most-exciting stimuli. To that end, we ranked neurons (with an aRF) from each field-of-view (FOV) following two criteria, equally weighted: (1) trial-to-trial reliability of the recorded responses, *i.e*. the pairwise correlation of the recorded responses to the same stimulus across all repeated presentations; (2) model prediction performance for the neuron, as measured by the trial-averaged correlation between predicted and recorded responses.

### 2.4 Most-exciting stimuli for neuronal populations

The procedure to find and generate most-exciting stimuli to neuronal populations is the same as the one used for single neurons (Methods 4.6) with two notable differences: (1) stimuli were centred using the population aRF, *i.e*. the aRFs average over all neurons in the population; (2) the predicted response was computed as the summed response over the stimulus presentation period, averaged over trials **and** neurons. Since the prediction performance of the model was not uniform across neurons, we computed the population average response weighted by the prediction performance of the model in the validation set.

### 4.8 *In vivo* two-photon calcium imaging of mouse V1 neurons (Rochefort laboratory)

#### 4.8.1 Animals

Animal experiments were approved by the Animal Welfare and Ethical Review Board (AWERB) of the University of Edinburgh and were performed under a project license granted by the UK Home Office conforming to the UK Animals (Scientific Procedures) Act 1986 and the European Directive 86/609/EEC and 2010/63/EU on the protection of animals used for experimental purposes. This study used 8-16 week old female mice on a C57BL/6J background (RRID: IMSR_JAX:010908). Mice were group housed in a reverse light/dark 12-hour cycle room that was kept at 21 ± 2°*C* and 55 ± 10% humidity, and had *ad libitum* access to food (DBM Scotland Ltd UK) and water.

#### 4.8.2 AAV injection and cranial window

Mice were anaesthetised during surgery using isoflurane and mounted on a stereotaxic frame (David Kopf Instruments, CA), with body temperature maintained at 37 °C using a servo-driven heater (Harvard Apparatus). Non-transparent eye cream (Bepanthen, Bayer, Germany) was applied to protect the eyes. The following analgesics and anti-inflammatory drugs were administered subcutaneously pre-operatively: buprenorphine (Veteregesic; 0.1 mg/kg), dexamethasone (Rapidexon; 2 mg/kg) and carprofen (Carprieve; 20 mg/kg). To prevent dehydration, an additional subcutaneous injection of warm Ringer’s saline solution (25 mL/kg; VWR, USA) was given at the end of the surgery. A section of the scalp was removed, and the underlying bone was cleared from tissue and blood. A single square craniotomy (2 × 2 mm) was made over the left primary visual cortex, centred at 2.5-3 mm lateral to midline and 1 mm anterior to lambda. After the craniotomy, adeno-associated (AAV) virus expressing the genetically encoded calcium indicator GCaMP (AAV1.Syn.GCaMP6f.WPRE.SV40; 10^10^ − 10^12^ IU/*µ*L; 1:5 in aCSF; UNC, Vector Core, Chapel Hill, NC) was injected using a micromanipulator and a pipette with a 20 µm tip diameter (Nanoject; Drummond Scientific, PA) at a speed of 10 nl/min, at three different depths at 60° angle (650, 550, and 450 µm from pia; 50-100 nl per site). Injections started at the deeper site. After each injection, the pipette was left *in situ* for an additional 5 min to prevent backflow. The craniotomy was then sealed with a custom-shaped glass coverslip (Menzel-Glaser #0) and fixed with cyano-acrylic glue. A custom-built round aluminium head-post was then implanted on the exposed skull with glue and secured in place with opaque dental acrylic (Paladur; Heraeus Kulzer, Germany). Mice were placed in a clean holding cage, positioned over a heating pad and monitored until they recovered from anaesthesia, before returning to their home cage. Imaging experiments started 3-6 weeks post-injection to allow for virus expression and clearing of the window.

#### 4.8.3 Habituation and head-fixation

Mice were extensively handled and habituated to head-fixation and the two-photon imaging set-up. Additionally, they were given *ad libitum* access to spinner exercise wheels in their homecage postsurgery.

#### 4.8.4 *In vivo* two-photon imaging

Mice were head-fixed onto a cylindrical polystyrene treadmill (20 cm diameter, on a ball bearing mounting axis) and allowed to run or sit freely. Imaging was performed using a custom-built resonant scanning two-photon microscope as described previously [34, 39, 58, 59]. In brief, the set up was equipped with a custom-built Ti:Sapphire excitation laser (Charmeleon Vision-S, Coherent, CA) tuned to 920 nm, and GaAsP photomultiplier tubes (Scientifica). Time-series images were acquired at a rate of 30 Hz using custom-programmed LabView based software (v8.2; National Instruments, UK) and a Nikon 25x water-immersion objective (Nikon; CF175 Apo 25XC W; 1.1 NA). Time-series images from 2 focal planes (field-of-view, FOV) per mouse were acquired on separate recording days at cortical depths between 160 µm to 260 µm from pia. Chronic imaging of the same FOVs was performed.

#### 4.8.5 Pupil imaging and recording of locomotion speed

Pupil dilation was monitored by recording the eye contralateral to the visual stimulation screen using a camera (monochrome CMOS DMK22UX273 camera; ImagingSource) coupled with a fixed focal length lens (Computar M5018-MP2), positioned ~15 cm from the mouse. Time-series images were acquired at a rate of 30 Hz using the IC Capture Image Acquisition software (ImagingSource).

Locomotion speed was measured with an optical encoder (2500 cpr, Pewatron, Switzerland) connected to a data acquisition device (National Instruments, UK) running LabView software (National Instruments, UK), sampled at 12 kHz and subsequently down-sampled to match the imaging sampling frequency (30 Hz).

#### 4.8.6 Visual stimuli

Visual stimuli were displayed on an LCD monitor (51 cm by 29 cm; Dell, UK) placed in front of the contralateral eye (right), 20 cm from the eye, using the Psychophysics Toolbox package [15] for MATLAB (MathWorks, MA). At the beginning of the first imaging session, we first performed brief retinotopic mapping to optimise the position of the screen which was kept constant between recording days.

##### Day 1 Natural movie viewing

The composition of the natural movie dataset follows the Sensorium dataset [81] (see Methods 4.2.1). 800 non-overlapping 10 s clips were randomly sampled from the following 8 movies and documentaries: Her (2013), Inception (2010), Jurassic Park (1993), Ratatouille (2007), Rogue One: A Star Wars Story (2016), The Matrix (1999), Tiny Titans - The Fascinating Lives of Rodents (2024) and Zootopia (2016). Each movie clip was downsampled to 1920 by 1080 pixels, converted to grey-scale and resampled to be displayed at 30 Hz. We then randomly split the movie clips into train, validation and test sets with a split ratio of 70%, 10% and 20%. For each FOV, we first randomly selected 410, 5 and 10 unique movie clips from the train, validation and test sets, respectively. We randomly ordered all 10 s movie clips and added a sequence of 0.5 s grey, 0.2 s white and 0.5 s grey screen between each clip. The training movies were presented once, while the validation and test movies were presented 10 times each. In total, we presented 560 natural movie clips for 103 min. We used OpenCV [14] to encode the videos into the AVI file format.

##### Day 2 Test of the model-generated stimuli and predictions

For the second session of each field-of-view (FOV), we first presented to the animal the same test set movies (10 unique movies, each with 5 repeats) in order to be able to test the session-to-session reliability of neuronal responses. We then presented artificial and model-generated stimuli to the animal in order to verify our model predictions (see Results 2.5, Methods 4.6 and Methods 4.7). Supp. Section S6 shows the stimuli that were presented to each mouse during the second session of each FOV. All stimuli were presented in the same manner as *in silico*, namely, each stimulus was presented for 1 s (30 frames) with 5 to 10 stimulus presentations. A grey screen of 1 s is added between each stimulus presentation. All artificial stimuli (low-*vs*. high-contrast centre-surround gratings, grating *vs*. natural *vs*. generated surround, etc.) were all combined to a single long video and presented in randomised order. To display on our computer screen the most-exciting stimuli, which were generated in 64 by 36 pixels (spatial resolution of the Sensorium dataset), we upscaled the stimuli to 1920 by 1080 pixels via bilinear interpolation with antialiasing (torchvision.transforms.functional.resize).

#### 4.8.7 Two-photon calcium imaging analysis

Image analysis for two-photon calcium imaging was performed as previously described [34, 39]. Briefly, following image acquisition at 30 Hz, brain motion was corrected using the toolbox NoRM-Corre [63]. Regions of interest (ROIs) corresponding to cell bodies were manually segmented by inspecting down-sampled frames (2 Hz), as well as the maximum intensity projection of each imaging stack, using ImageJ/FIJI software [70] (NIH public domain; RRID: SCR-002285). To match ROIs across days, we used a semi-automatic method based on spatial pairwise distance and angular similarity scoring of ROI centroids (github.com/rochefort-lab/ROI_matching). Pixel fluorescence within each ROI was averaged to create a raw time series *F* (*t*). The baseline fluorescence (*F*_0_) was computed by taking the 5^th^ percentile of the smoothed *F(t)* (1 Hz low-pass, zero-phase, 60^th^-order FIR filter) per ROI over each trial (*F*_0_(*t*)), averaged across all trials. The Δ*F/F*_0_ was then computed by taking the difference between *F* (*t*) and *F*_0_(*t*) and dividing it by *F*_0_. To remove neuropil contamination, we used the toolbox Fast Image Signal Separation Analysis (*FISSA*) [40]. Python toolboxes (*SIMA* and *FISSA*) were run with WinPython 2.7.10.3 and further analysis was performed with custom-written scripts in MATLAB (MathWorks, MA) and Python 3.12.7.

#### 4.8.8 Analysis of pupil size and position

Pupil diameter was measured from the videos of the eye using DeepLabCut [50]. For each animal 180 training frames were randomly sampled from the trials. Six points were manually identified and spaced approximately evenly around the pupil. The network (`resnet-50`) was trained with default parameters for a maximum of 100k to 200k iterations. The pupil size was computed as the area of the smallest enclosing circle to the detected pupil points. If points were labelled with a likelihood of smaller than 0.9 (*e.g*., blinks), they were replaced with nearest-neighbour interpolation.

#### 4.8.9 Training ViV1T with new *in vivo* imaging dataset

We tested three different methods to (re)train ViV1T on the *in vivo* recordings: direct, transfer and fine-tune (see Results 2.2), using the same hyperparameters from Methods 4.2.2. To stay consistent with the model trained on the Sensorium dataset, we also downsampled the videos to a resolution of 36 height by 64 width pixels. Given its superior prediction performance, we used the fine-tuned models in all analyses of the *in vivo* imaging data from Rochefort laboratory.

## Acknowledgments

We sincerely thank Turishcheva *et al*. for making their high-quality large-scale mouse V1 neuronal recordings publicly available thereby making this work possible. We thank Dr. Matthias H. Hennig and Dr. Nina Kudryashova, Dr Theoklitos Amvrosiadis and all members of the Rochefort laboratory for their insightful feedback on early versions of the manuscript. We thank the GENIE Program and the Janelia Research Campus, specifically V. Jayaraman, R. Kerr, D. Kim, L. Looger, and K. Svoboda, for making GCaMP6 available. We would like to express our gratitude to the dedicated staff at Bioresearch and Veterinary Services (BVS) at the University of Edinburgh for their invaluable support throughout our animal work. BML and WDW were supported by the United Kingdom Research and Innovation (grant EP/S02431X/1), UKRI Centre for Doctoral Training in Biomedical AI at the University of Edinburgh, School of Informatics. This work was funded by the Simons Initiative for the Developing Brain (to D.K. and N.L.R.), the European Research Council (grant agreement 866386 to N.L.R.), the Shirley Foundation, the Patrick Wild Centre, the European Molecular Biology Organisation (YIP award to N.L.R.). This work was supported by the Edinburgh International Data Facility (EIDF) and the Data-Driven Innovation Programme at the University of Edinburgh.

## Author contributions

We use the contribution categories from CRediT (Contributor Roles Taxonomy). Authors within each category are sorted in the same order as in the author list. Conceptualization: BML, AO and NLR. Data Curation: BML and DK. Formal analysis: BML, WDW and DK. Funding acquisition: NLR and AO. Investigation: BML, WDW and DK. Methodology: BML, WDW and AO. Project administration: AO and NLR. Resources: BML, WDW and AO. Software: BML and WDW. Supervision: AO and NLR. Validation: BML and WDW. Visualisation: BML. Writing – original draft: BML, WDW, DK, AO and NLR. Writing – review & editing: BML, WDW, DK, AO and NLR.

## Supplementary Material

### S1 Additional prediction performance assessment

In addition to the normalized correlation (Methods 4.3), we also evaluated predictive performance of models using the single trial Pearson correlation *r*_trial_ between recorded and predicted responses, as used in the Sensorium competition [81]:

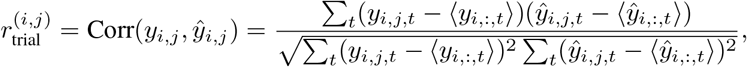

where *y*_*i,j,t*_ and *ŷ*_*i,j,t*_ denote recorded and predicted responses, respectively, for neuron *i* on trial *j* at time *t*, and ⟨·⟩ here denotes average across trials/stimulus repeats. Like in the Sensorium competition, *r*_trial_ is calculated independently per neuron and trial, and then averaged across all neurons and trials to obtain a population metric.

**Supplemental Table 1.** Single trial correlation (s.e.m. across animals) between recorded and predicted responses in movie and classical stimuli test sets. The most and second most performing model in each category highlighted in **bold** and <u>underline</u>, respectively.

| Model | Natural movie | Pink noise | Flashing dot | Moving dots | Drifting Gabor | Image | Avg. |
| --- | --- | --- | --- | --- | --- | --- | --- |
| LN | 0.112<br>(0.002) | 0.057<br>(0.002) | 0.081<br>(0.004) | 0.045<br>(0.006) | 0.078<br>(0.005) | 0.115<br>(0.004) | 0.081 |
| fCNN | 0.246<br>(0.005) | 0.107<br>(0.005) | 0.121<br>(0.006) | 0.109<br>(0.011) | 0.127<br>(0.003) | 0.215<br>(0.011) | 0.154 |
| DwiseNeuro | <u>0.284</u><br>(0.007) | <u>0.151</u><br>(0.004) | **0.162**<br>(0.009) | <u>0.147</u><br>(0.010) | **0.188**<br>(0.005) | <u>0.255</u><br>(0.012) | <u>0.198</u> |
| ViVIT | **0.298**<br>(0.008) | **0.161**<br>(0.006) | <u>0.156</u><br>(0.009) | **0.152**<br>(0.013) | <u>0.187</u><br>(0.002) | **0.269**<br>(0.017) | **0.204** |
| ViVIT<br>(without behaviour) | 0.255<br>(0.010) | 0.126<br>(0.001) | 0.108<br>(0.011) | 0.110<br>(0.007) | 0.139<br>(0.009) | 0.219<br>(0.013) | 0.160 |

**Supplemental Table 2.** Trial-averaged correlation (s.e.m. across animals) between recorded and predicted responses in movie and classical stimuli test sets. The most and second most performing model in each category highlighted in **bold** and underline, respectively.

| Model | Natural movie | Pink noise | Flashing dot | Moving dots | Drifting Gabor | Image | Avg. |
| --- | --- | --- | --- | --- | --- | --- | --- |
| LN | 0.142<br>(0.008) | 0.090<br>(0.001) | 0.138<br>(0.012) | 0.046<br>(0.001) | 0.110<br>(0.006) | 0.171<br>(0.007) | 0.116 |
| fCNN | 0.305<br>(0.005) | 0.196<br>(0.002) | 0.217<br>(0.012) | 0.189<br>(0.012) | 0.198<br>(0.009) | 0.325<br>(0.013) | 0.238 |
| DwiseNeuro | <u>0.363</u><br>(0.008) | 0.277<br>(0.003) | <u>0.284</u><br>(0.020) | <u>0.255</u><br>(0.011) | **0.287**<br>(0.013) | <u>0.382</u><br>(0.018) | <u>0.308</u> |
| ViVIT | **0.382**<br>(0.008) | **0.297**<br>(0.006) | **0.285**<br>(0.023) | **0.269**<br>(0.011) | <u>0.279</u><br>(0.010) | **0.403**<br>(0.019) | **0.319** |
| ViVIT<br>(without behaviour) | 0.353<br>(0.008) | <u>0.278</u><br>(0.006) | 0.232<br>(0.018) | 0.241<br>(0.011) | 0.252<br>(0.013) | 0.369<br>(0.015) | 0.287 |

### S2 Prediction of spatial organisation of orientation selective neurons

**Supplemental Figure 1.**
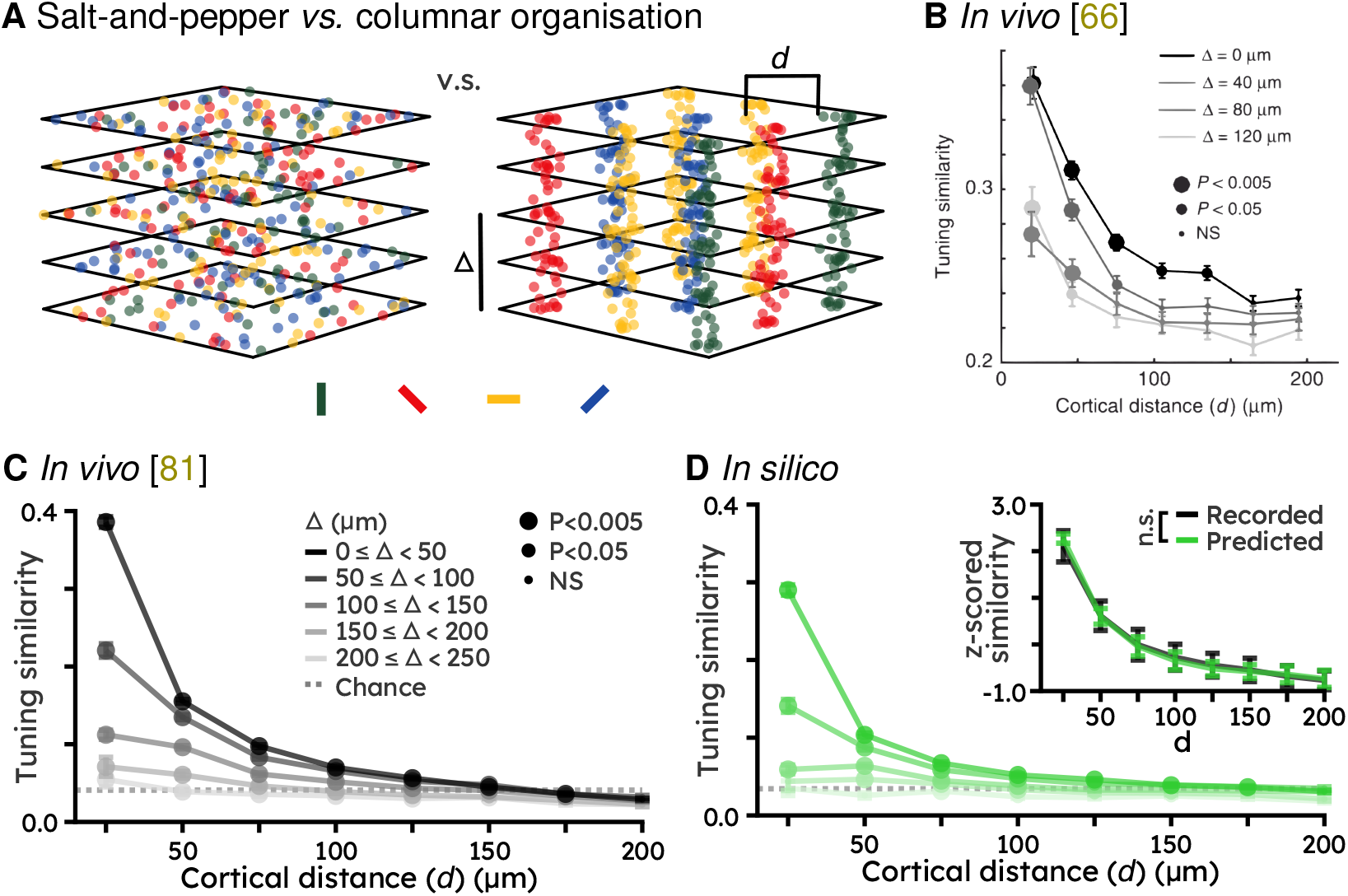
Movie-trained ViV1T predicts spatial clustering of mouse V1. (A) Schematic representation of salt-and-pepper (left) and columnar (right) organisation of orientation selective neurons across V1 superficial layers. *d* indicates within-plane cortical distance, and Δ across-plane depth distance. (B) Ringach *et al*. [66] show that the orientation tuning similarity of pairs of mouse V1 neurons within or across different cortical planes decays as a function of *d* and Δ. For each individual curve, the size of the data points denotes the significance of a rank-sum test comparing the median distribution of data at a given distance to the distribution of the rightmost bin. Error bars indicate s.e.m. over neurons, pooled across 4 mice. Reproduced from Ringach *et al*. [66]. (C) Same as (B), but for the Sensorium dataset. Error bars indicate s.e.m. over neurons, pooled across mice (*N*_neurons_ = 4095 from *N*_mice_ = 3). Dotted line shows the average tuning similarity of 5,000 randomly selected neuron pairs. (D) Same as (C), but using ViV1T model predictions. Inset shows curves averaged across Δ. Error bars indicate s.e.m. over mice. Log-likelihood ratio test finds individual exponential decay fits not significantly better than a common fit (ratio = 5.941, n.s.: *p* = 2.036 × 10^−1^), see Methods 4.4 for details.

### S3 Comparison of ViV1T predictions against other state-of-the-art models

We showed that ViV1T performed better than other models at predicting V1 responses to full-field natural movies and artificial stimuli (Table 1). We also compared ViV1T to other models in their ability to predict V1 neuronal tuning properties and responses to contextual modulation. Altogether, ViV1T performed either better or as well as other models across the different visual response properties that were tested. In addition, our model is significantly faster to run (20 times faster) and requires fewer parameters (13 times smaller) than the second best performing model (see also Results 2.1).

**Supplemental Table 3.**
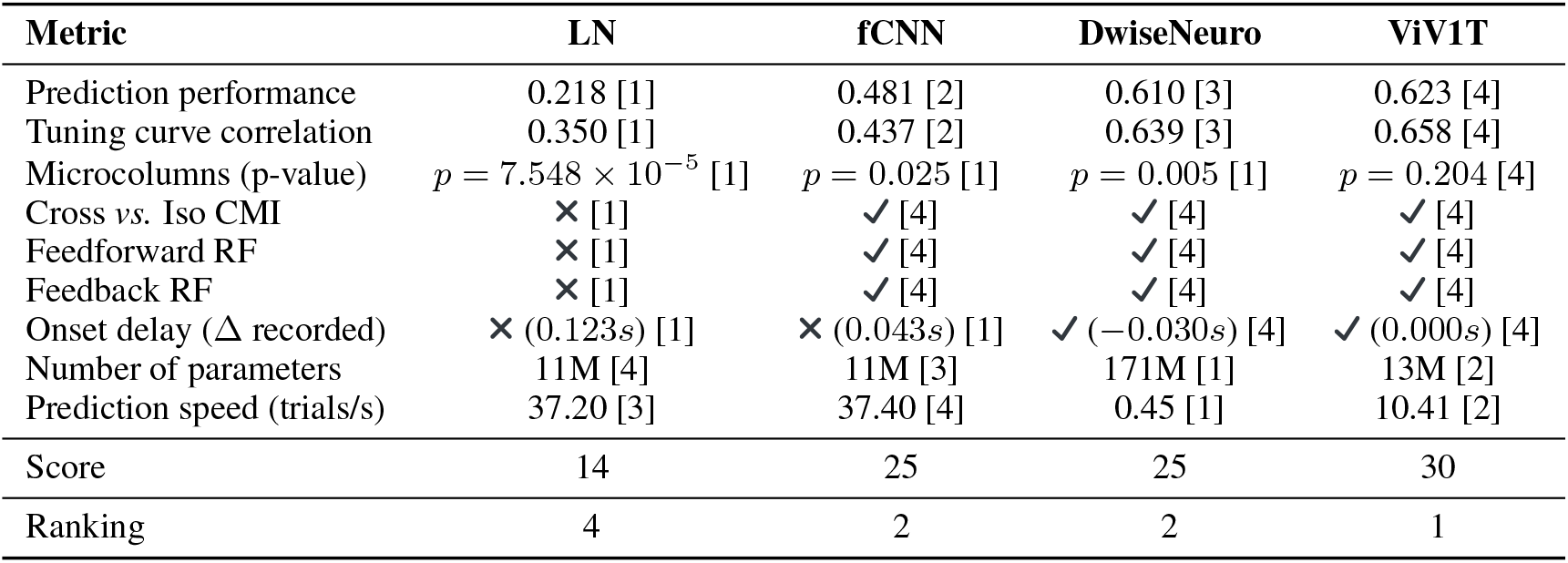
Model comparison across V1 neuronal properties. For metrics with continuous values, we rank models from best to worst, assigning scores from 4 to 1, shown in square brackets. Binary metrics grant either score 4 (predicted) or score 1 (not predicted). We add up all scores and rank the models accordingly.

**Supplemental Figure 2.**
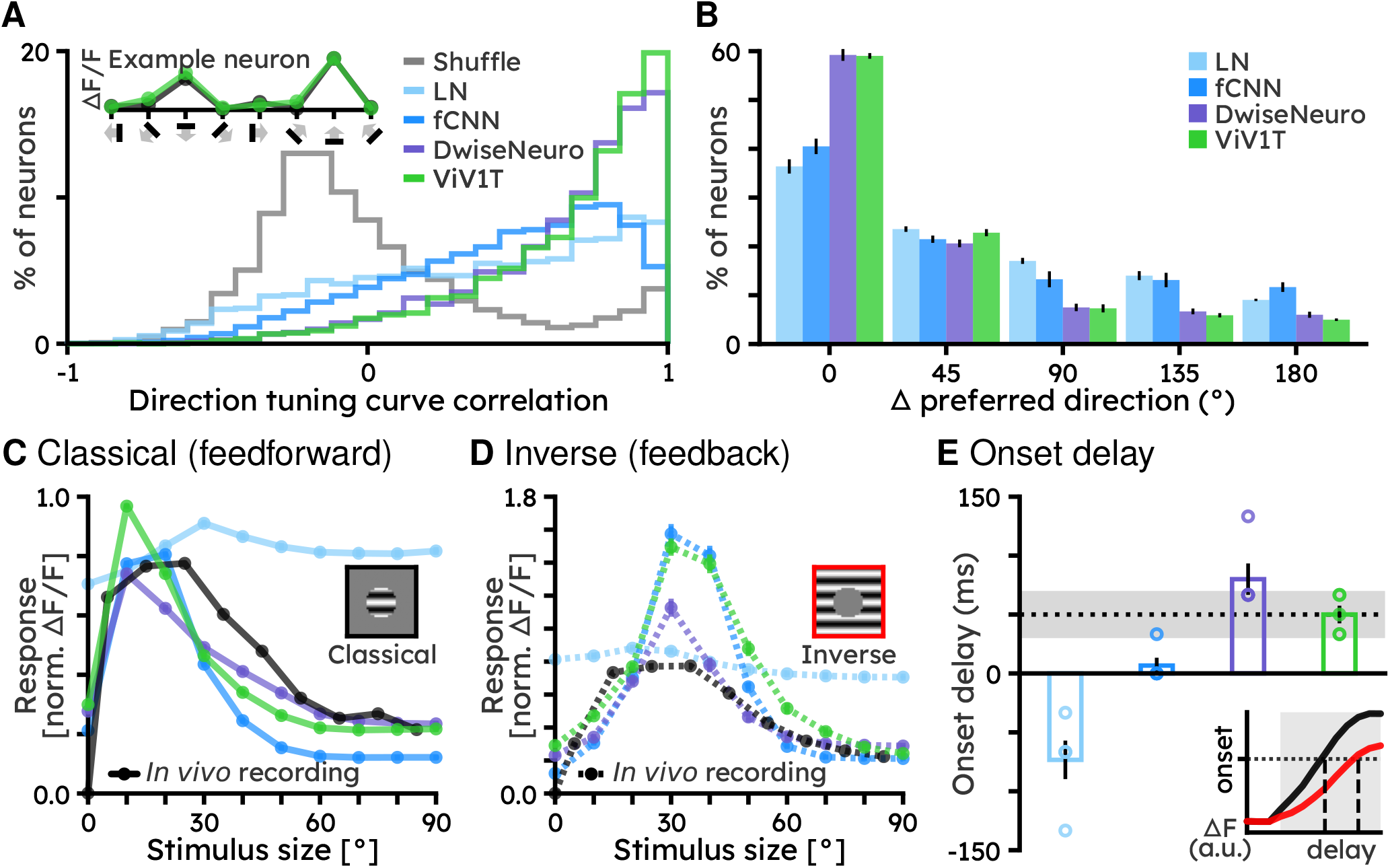
Model comparison for direction tuning, and feedforward/feedback receptive fields. (A) Histogram of correlation values between recorded and predicted direction tuning curves for all models (coloured) and shuffled recorded tuning curves (grey). Inset shows recorded (black) and ViV1T predicted (green) direction tuning curves of example neuron 1. (B) Difference between recorded *vs*. predicted preferred direction for each of the models. (C) and (D) Model-predicted population-averaged size tuning curves to classical and inverse stimuli, normalised to the maximum response for classical stimuli. Recorded results from Keller *et al*. [42] are presented in black. (E) Difference between inverse and classical response onsets. Dotted line illustrates the average response onset delay of 50±20 ms recorded *in vivo* [42]. Inset illustrates delay between (red) inverse and (black) classical response onsets.

### S4 ViV1T predictions of contextual responses for low vs high contrasts in the Sensorium dataset

**Supplemental Figure 3.**
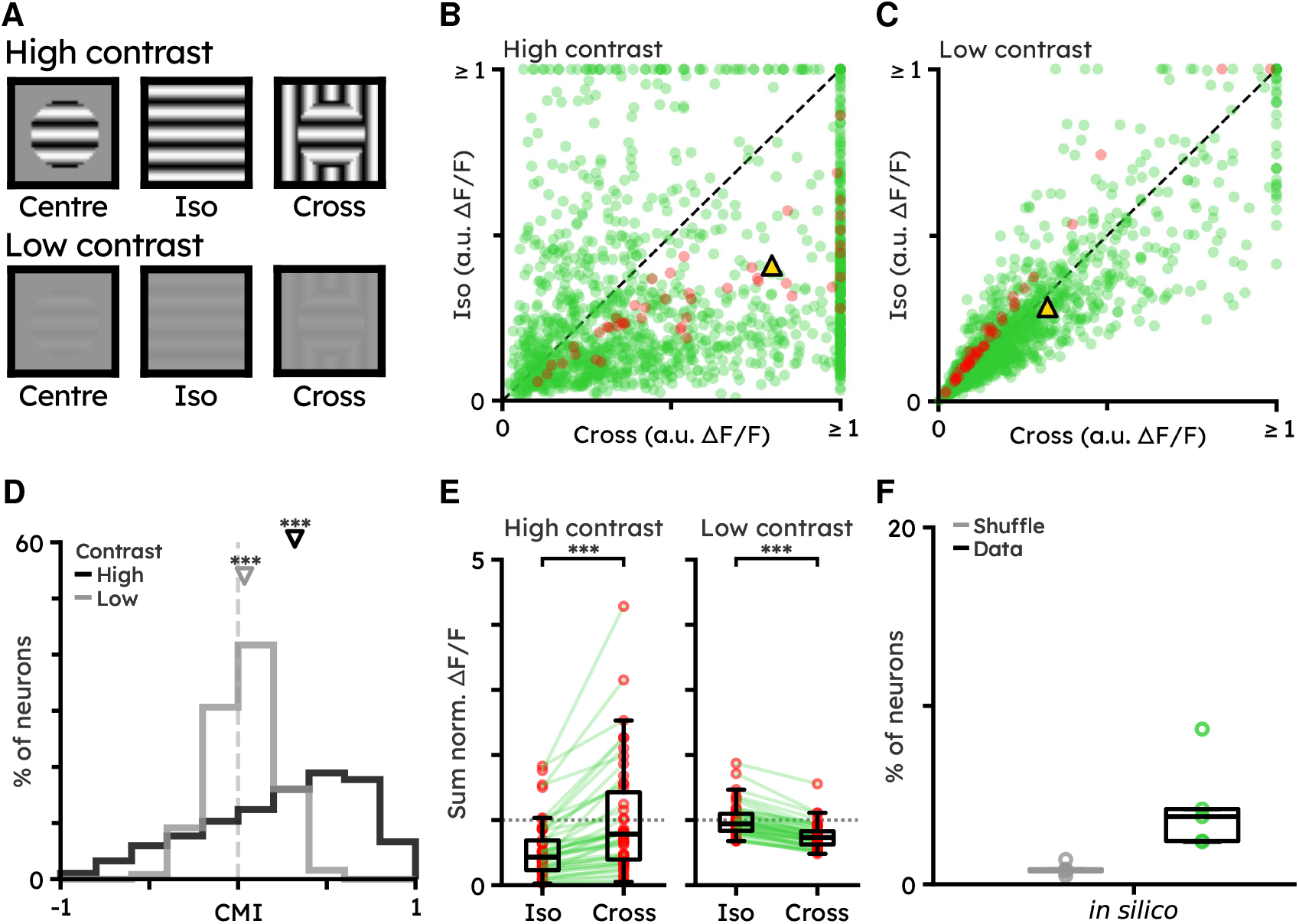
A subpopulation of V1 neurons switches contextual modulation with contrast in the Sensorium dataset. (A) Illustration of the centre, iso-surround and cross-surround Gabor grating stimuli in high (top row) and low (bottom row) contrast. (B) *In silico* responses to high contrast iso- and cross-surround gratings. Each dot is the trial-averaged Δ*F/F* for each neuron. Gold triangle indicates the population average. Red dots show neurons that switch contextual modulation with contrast. (two-sided Wilcoxon signed-rank test: *N*_neurons_ = 1443 from *N*_mice_ = 5, *p <* 1 × 10^−10^). (C) Same as (B) but with low contrast stimulus (two-sided Wilcoxon signed-rank test: *p <* 1 × 10^−10^). (D) Contextual modulation index (high contrast, mean: 0.243, (black triangle) median: 0.320; low contrast, mean: 0.035, (grey triangle) median: 0.036; two-sided Wilcoxon signed-rank test, high contrast ***: *p <* 1 × 10^−10^, low contrast ***: *p <* 1 × 10^−10^). (E) *In silico* neurons that switch contextual modulation between high- and low-contrast stimuli. All responses are normalised by response to centre grating at high contrast (dotted line). (two-sided Wilcoxon signed-rank test: *N*_neurons_ = 46, high contrast ***: *p <* 1 × 10^−10^, low contrast ***: *p <* 1 × 10^−10^). (F) Percentage of (green) *in silico* neurons that switch contextual modulation. Each dot indicates the percentage of neurons switching contextual modulation between high and low contrast stimuli per mouse. Grey boxes indicate the results obtained from shuffled responses.

### S5 ViV1T accurately predicts V1 population neuronal responses

**Supplemental Figure 4.**
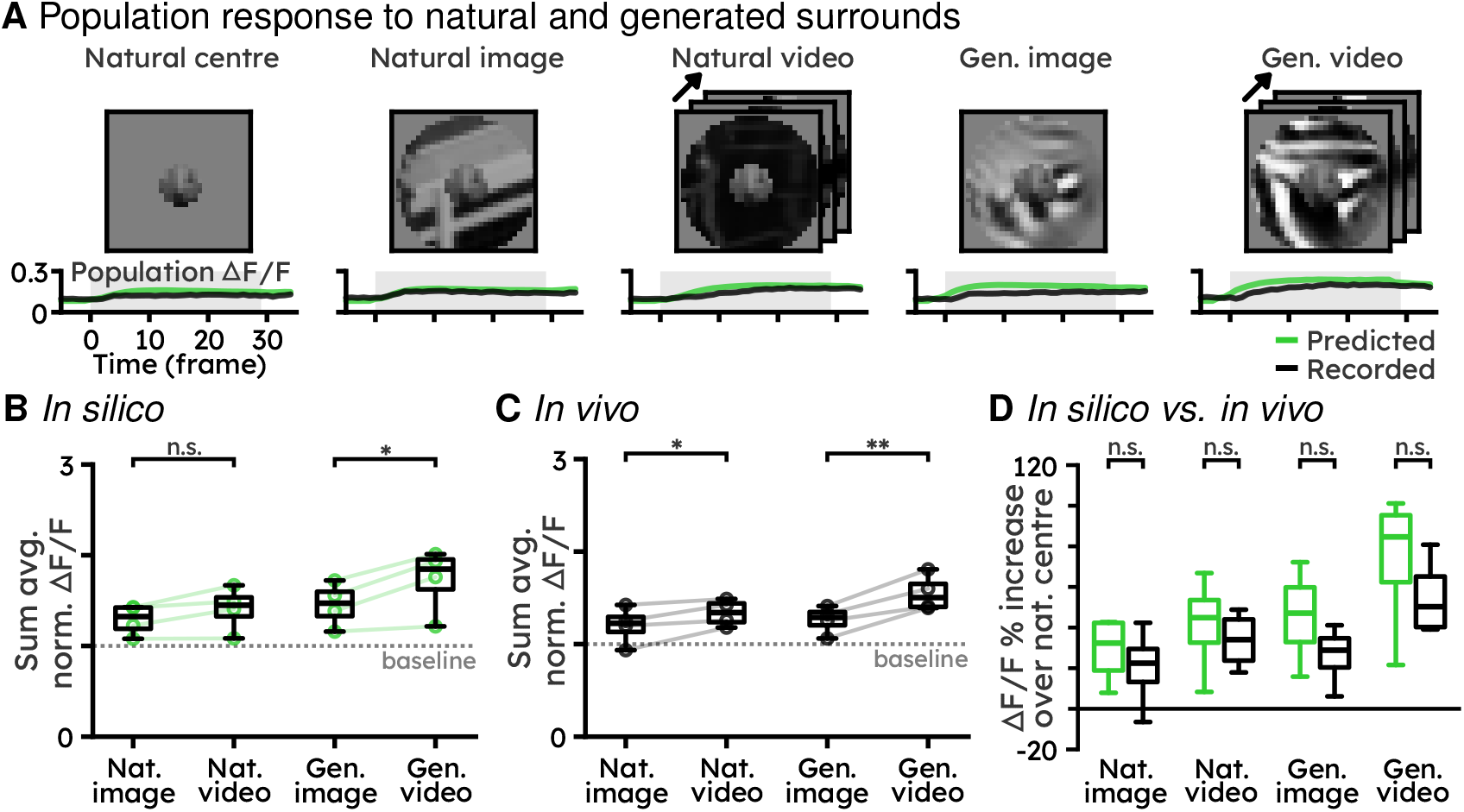
ViV1T accurately predicts surround modulation of V1 population neuronal responses. (A) Example of contextual responses of all V1 imaged neurons in one field-of-view. The left-most column shows the average response across all neurons to the most-exciting natural image, centred on the population’s receptive field. The right columns show the responses of the same neuronal population to stimuli combining the same centre with the most-exciting natural image, natural video and ViV1T-generated image and video surrounds. The black and green lines are the trial-averaged recorded and predicted population Δ*F/F*. (B) *In silico* average population responses normalised by the response to the most exciting natural centre stimulus. Individual circles are FOVs, error bars indicate s.e.m. over FOVs (one-sided t-test: *N*_FOVs_ = 4; natural image *vs*. natural video n.s.: *p* = 0.052; generated image *vs*. generated video *: *p* = 0.018). (C) Same as (B) but for *in vivo* recordings. Both natural and model-generated dynamic stimuli evoked larger responses than their static counterparts (B), with a median increase of 10% and 21%, respectively (One-sided t-test: *N*_FOVs_ = 4; natural image *vs*. natural video *: *p* = 0.028; generated image *vs*. generated video **: *p* = 0.006). (D) Population response increase over baseline (most-exciting natural image centre stimuli) for (green) *in silico* prediction and (black) *in vivo* recordings (two-sided t-test between *in silico* and *in vivo*: natural image n.s.: *p* = 0.619; natural video n.s.: *p* = 0.574; generated image n.s.: *p* = 0.340; generated video n.s.: *p* = 0.403).

### S6 Additional dataset information

#### Sensorium 2023 dataset

**Supplemental Table 4.** Test set labels (*in vivo* recordings) availability in the Sensorium dataset [81].

| Mouse | ID | $N_{\text{neurons}}$ | Natural movie | Pink noise | Gaussian dots | RDK | Drifting Gabor | Image |
| --- | --- | --- | --- | --- | --- | --- | --- | --- |
| A | dynamic29156-11-10 | 7440 | ✓ | ✓ | ✓ | ✓ | × | × |
| B | dynamic29228-2-10 | 7928 | ✓ | ✓ | × | × | ✓ | ✓ |
| C | dynamic29234-6-9 | 8285 | ✓ | × | × | ✓ | ✓ | ✓ |
| D | dynamic29513-3-5 | 7671 | ✓ | ✓ | ✓ | × | × | ✓ |
| E | dynamic29514-2-9 | 7495 | ✓ | × | ✓ | ✓ | ✓ | × |
| F | dynamic29515-10-12 | 7863 | × | × | × | × | × | × |
| G | dynamic29623-4-9 | 7908 | × | × | × | × | × | × |
| H | dynamic29647-19-8 | 8202 | × | × | × | × | × | × |
| I | dynamic29712-5-9 | 7939 | × | × | × | × | × | × |
| J | dynamic29755-2-8 | 8122 | × | × | × | × | × | × |

#### Rochefort Lab dataset

**Supplemental Table 5.** Rochefort Lab *in vivo* recording composition. In total, *in vivo* recordings from 2 mice were obtained, each from 2 different field-of-views (FOVs). On the first session of recording (Day 1), about 2 h of natural movies were presented to the animals which is used to train the ViV1T model (see Methods 4.8.9). We then showed the low-vs. high-contrast centre-surround gratings, model-selected and model-generated stimuli in the second session(s). Neurons in recorded in day 2 and 3 were matched against those recorded on day 1. Natural movies were presented to all FOVs on day 1. A subset of the natural movies presented on day 1 were presented in the beginning of the day 2 (and 3, if exists) recording session so that we can measure the session-to-session reliability.

| Mouse | FOV | $N_{\text{neurons}}$ | | | Stimuli presented | | | |
| --- | --- | --- | --- | --- | --- | --- | --- | --- |
|  |  | Day 1 | Day 2 | Day 3 | Fig. 4 | Figs. 5A to 5D | Figs. 5E to 5H | Supp. Figure 4 |
| 232 | K | 350 | 212 | NA | ✓ | × | ✓ | ✓ |
| 232 | L | 216 | 146 | 133 | ✓ | ✓ | ✓ | ✓ |
| 233 | M | 352 | 245 | NA | ✓ | ✓ | × | ✓ |
| 233 | N | 252 | 205 | NA | ✓ | ✓ | ✓ | ✓ |

### S7 Trade off between ViV1T performance and number of stimuli and neurons used for model training

The Sensorium dataset is a large-scale dataset acquired for a machine learning competition [81, 82]. It consists of two-photon calcium imaging of around 8,000 V1 neurons per mouse from 10 mice, during the presentation of hundreds (about 350) of 10 s action movie clips. However, not all experimental laboratories are capable of recording such large datasets. Moreover, as demonstrated with our own *in vivo* recordings (Methods 4.8 and Methods 4.8.9), it may not be necessary to record that many neurons to test a given hypothesis. We systematically investigated how much training data and neurons are needed to train ViV1T to reach different levels of performance (accurate predictions of visual responses to natural movies) Mineault *et al*. [53].

We trained ViV1T on natural movies, with 80 different combinations of numbers of training samples (number of movie clips) and neurons from the Sensorium dataset. We then compared the prediction performance for each combination to unseen natural movies and artificial stimuli (drifting gratings, natural images, pink noise, flashing dots, moving dots) against the model that was trained on all training data and neurons (Table 1). Supp. Figure 5 shows the prediction performance of each model configuration as a percentage of the performance of the full model. As expected, prediction performance improved with more training samples and neurons, with training samples (number of movie clips) being the larger limiting factor. A prediction performance above 85% (*i.e*. normalised correlation of 0.592) of maximum performance with the full model was reached with 100 neurons and 90% of the training samples (about 315 movie clips). A similar trend was observed for predictions of responses to artificial stimuli with the number of neurons having more impact (Supp. Figure 5).

**Supplemental Figure 5.**
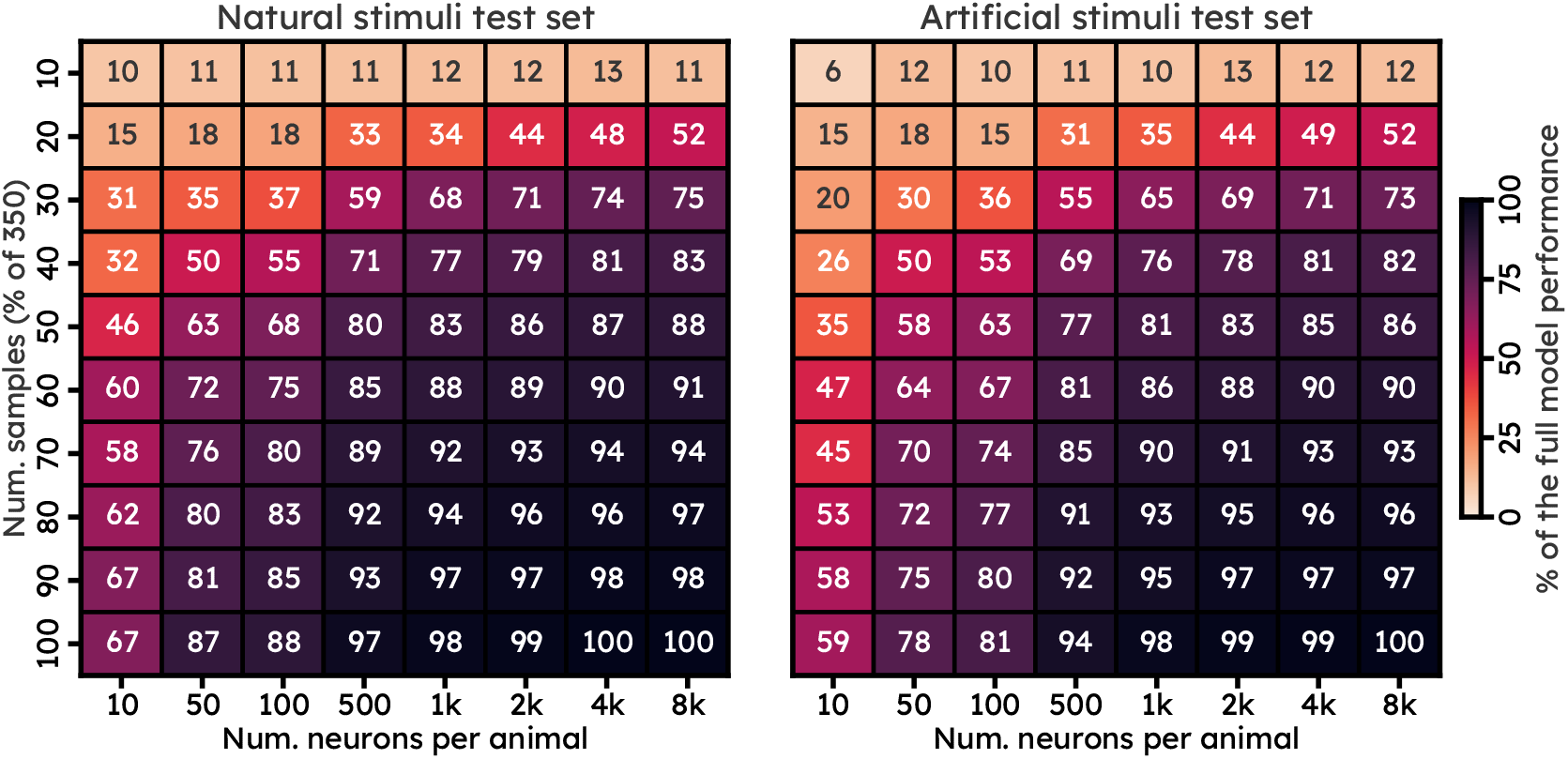
Prediction performance of ViV1T trained with varying numbers of training samples and neurons per animal. Predictions of neuronal responses for unseen natural movies are shown on the left and for artificial stimuli (drifting gratings, flashing images, flashing dots, pink noise and moving dots). Results are presented as a percentage of the prediction performance of the full model (*i.e*. Table 1) which was trained on all training data (350 movie clips) and neurons (8000 neurons per animal from 10 mice). The training samples and neurons in each row and column are the same, *i.e*. the same 10 neurons from each animal were used in training all model combinations presented in the first column. Neurons and training samples were randomly subsampled via `np.random.choice`with no repeat.

### S8 Hyperparameters

**Supplemental Table 6.** ViV1T hyperparameter search space and their final settings after a Hyperband Bayesian optimization [46].

| HYPERPARAMETER | SEARCH SPACE | FINAL VALUE |
| --- | --- | --- |
| **CORE** |  |  |
| NUM. SPATIAL BLOCKS | UNIFORM, MIN: 1, MAX: 16 | 14 |
| NUM. TEMPORAL BLOCKS | UNIFORM, MIN: 1, MAX: 16 | 14 |
| NUM. HEADS | UNIFORM, MIN: 1, MAX: 12 | 2 |
| SPATIAL PATCH SIZE | UNIFORM, MIN: 3, MAX: 9 | 7 |
| SPATIAL PATCH STRIDE | UNIFORM, MIN: 1, MAX: SPATIAL PATCH SIZE | 2 |
| TEMPORAL PATCH SIZE | UNIFORM, MIN: 1, MAX: 50 | 13 |
| PATCH METHOD | SLIDING WINDOW, 2D CONV, SPT, CCT | SLI. WIND. |
| PATCH DROPOUT | UNIFORM, MIN: 0, MAX: 0.5 | 0.17 |
| EMBEDDING SIZE | UNIFORM, MIN: 64, MAX: 256, INTERVAL: 8 | 112 |
| HEAD SIZE | UNIFORM, MIN: 8, MAX: 256, INTERVAL: 8 | 80 |
| MHA DROPOUT | UNIFORM, MIN: 0, MAX: 0.5 | 0.04 |
| MLP SIZE | UNIFORM, MIN: 64, MAX: 256, INTERVAL: 8 | 224 |
| MLP ACTIVATION | TANH, ELU, ReLU, GELU, SiLU, SwiGLU | ReLU |
| MLP DROPOUT | UNIFORM, MIN: 0, MAX: 0.5 | 0.21 |
| NORMALIZATION | LAYER NORM, RMS NORM | LAYER NORM |
| REGISTRY TOKENS | UNIFORM, MIN: 0, MAX: 32 | 24 |
| STOCHASTIC DEPTH DROPOUT | UNIFORM, MIN: 0, MAX: 0.5 | 0.43 |
| WEIGHT DECAY | UNIFORM, MIN: 0, MAX: 1 | 0.0783 |
| CORE LEARNING RATE | UNIFORM, MIN: 0.0001, MAX: 0.1 | 0.0126 |
| **READOUT** |  |  |
| POSITION NETWORK NUM. LAYERS | UNIFORM, MIN: 1, MAX: 4, INTERVAL: 1 | 1 |
| POSITION NETWORK NUM. UNITS | UNIFORM, MIN: 2, MAX: 128, INTERVAL: 2 | 30 |
| BIAS INITIALIZATION | 0, MEAN STANDARDIZED RESPONSE | 0 |
| WEIGHT DECAY | UNIFORM, MIN: 0, MAX: 1 | 0.6893 |
| LEARNING RATE | UNIFORM, MIN: 0.0001, MAX: 0.1 | 0.0073 |
| OUTPUT ACTIVATION | NONE, ELU + 1, EXPONENTIAL, SOFTPLUS, SIGMOID | ELU + 1 |

**Supplemental Table 7.** Top 20 ViV1T hyperparameter importance in Hyberhand Bayesian Optimization [46] via Weights & Biases [13]. IMPORTANCE shows the degree to which the hyperparameter is useful to predict the evaluation metric (e.g. single trial correlation in the validation set) and CORRELATION shows the linear correlation between the hyperparameter and the evaluation metric. Details on the calculation and interpretation of the hyperparameter importance and correlation are available at docs.wandb.ai/guides/app/features/panels/parameter-importance.

| HYPERPARAMETER | IMPORTANCE | CORRELATION |
| --- | --- | --- |
| CORE LEARNING RATE | 0.138 | -0.236 |
| PATCH DROPOUT | 0.134 | -0.300 |
| CORE WEIGHT DECAY | 0.077 | -0.132 |
| EMBEDDING SIZE | 0.066 | -0.075 |
| READOUT LEARNING RATE | 0.061 | -0.042 |
| READOUT WEIGHT DECAY | 0.053 | 0.094 |
| STOCHASTIC DEPTH DROPOUT | 0.051 | -0.014 |
| NUM. TEMPORAL BLOCKS | 0.046 | -0.040 |
| NUM. SPATIAL BLOCKS | 0.042 | 0.083 |
| REGISTRY TOKENS | 0.042 | 0.049 |
| SPATIAL PATCH SIZE | 0.037 | -0.055 |
| TEMPORAL PATCH SIZE | 0.036 | -0.022 |
| MLP SIZE | 0.032 | -0.033 |
| NUM. HEADS | 0.029 | 0.34 |
| WARM UP STEPS | 0.029 | -0.14 |
| HEAD SIZE | 0.029 | 0.027 |
| MHA DROPOUT | 0.028 | 0.035 |
| MLP DROPOUT | 0.027 | 0.087 |
| OUTPUT ACTIVATION | 0.024 | -0.084 |
| SPATIAL PATCH STRIDE | 0.019 | 0.096 |

